# NRBP1 and TSC22D proteins impact distal convoluted tubule physiology through modulation of the WNK pathway

**DOI:** 10.1101/2024.12.12.628222

**Authors:** Germán Magaña-Ávila, Héctor Carbajal-Contreras, Ramchandra Amnekar, Toby Dite, Michelle Téllez-Sutterlin, Kevin García-Ávila, Brenda Marquina-Castillo, Alejandro Lopez-Saavedra, Norma Vazquez, Eréndira Rojas-Ortega, Eric Delpire, David H. Ellison, Dario R. Alessi, Gerardo Gamba, María Castañeda-Bueno

## Abstract

The With No lysine (WNK) kinases regulate processes such as cell volume and epithelial ion transport through the modulation of Cation Chloride Cotransporters such as the NaCl cotransporter, NCC, present in the distal convoluted tubule (DCT) of the kidney. Recently, the interaction of WNKs with Nuclear Receptor Binding Protein 1 (NRBP1) and Transforming Growth Factor β-Stimulated Clone 22 Domain (TSC22D) proteins was reported. Here we explored the effect of NRBP1 and TSC22Ds on WNK signaling in vitro and in the DCT. TSC22D1.1, TSC22D2, and NRBP1 are localized in DCT WNK bodies, which are cytoplasmic biomolecular condensates associated with WNK activation. In HEK293 cells, long TSC22D isoforms and NRBP1 increase WNK4 activity. DCT-specific NRBP1 knockout mice have reduced NCC phosphorylation and activate a compensatory response. Thus, NRBP1 and long TSC22D proteins are positive modulators of WNK signaling and modulate Na^+^ reabsorption in the kidney. NRBP1 and TSC22Ds likely influence WNK signaling in other tissues, impacting various physiological processes.

**Teaser:** The pseudokinase NRBP1 and its associated TSC22D proteins modulate WNK kinases to regulate sodium reabsorption in the kidney.

## Introduction

The With No lysine (K) (WNK) kinase family is comprised of four members (WNK1-4). These proteins have been implicated in the regulation of cell volume, epithelial electrolyte transport, neuronal intracellular chloride concentration, and cell proliferation (*1*). Most of these functions involve phosphorylation by WNK kinases of their canonical downstream targets Ste20-related Proline-Alanine-Rich Kinase (SPAK) and Oxidative Stress Response 1 (OSR1), that in turn phosphorylate and modulate the activity of Cation Chloride cotransporters (CCCs) of the Slc12 family. Among these CCCs, we find the loop diuretic-sensitive Na^+^:K^+^:Cl^-^ cotransporter (NKCC2) and the thiazide-sensitive Na^+^:Cl^-^ cotransporter (NCC), that mediate NaCl reabsorption in the thick ascending limb and distal convoluted tubule (DCT) of the kidney nephron, respectively.

Mutations in the genes encoding WNK1 and WNK4 are cause of a tubulopathy called Familial Hyperkalemic Hypertension (FHHt) (also known as Pseudohypoaldosteronism type II or Gordon syndrome) (*2*–*4*). Patients with FHHt present with hypertension, hyperkalemia, and metabolic acidosis. These electrolytic derangements are essentially due to the overactivity of NCC that is exclusively present in the apical membrane of the DCT (*5*, *6*). Thus, the role of WNK kinases in DCT physiology has been extensively studied. Evidence suggests that WNK4 is the major catalytically active WNK isoform in these cells (*7–9*). Moreover, in DCT the vast majority of WNK1 mRNA encodes a short isoform (kidney specific (KS)-WNK1) lacking kinase catalytic domain (*10*). Consistent with this, NCC activity and phosphorylation in the DCT, is completely lost in WNK4 knockout mice (*7*).

We and others have recently described that during osmotic stress, a condition in which WNK kinases are activated to promote ion influx to the cell through modulation of CCCs, the interaction of WNK1 with Nuclear Receptor Binding Protein 1 (NRBP1) and proteins of the Transforming Growth factor β-Stimulated Clone 22 Domain (TSC22D) family (*11*, *12*) is stimulated. The latter are known interactors of NRBP1 (*13*). The NRBP family is comprised of two members (NRBP1 and NRBP2) that are highly homologous. Interestingly, these proteins are pseudokinases that are evolutionarily related to WNK kinases (*12*, *14*). They present a general architecture similar to that of WNK kinases, with a short N-terminal domain, followed by the pseudokinase domain, and a C-terminal domain that is much smaller than that of WNK kinases, but that at least contains one common feature, which is a globular Conserved C-Terminal (CCT) domain (*12*) (also known as PF2 domain (*15–17*)). Similar CCT domains are present in WNKs and SPAK/OSR1. The CCT domains of SPAK and OSR1 have been shown to mediate the interaction of these kinases with RFxV motifs present in interacting partners like CCCs and WNKs. In addition, other protein binding motifs have been shown to be present in the C-terminal domain of NRBP1, like a Myeloid Leukemia Factor 1 binding motif (*18*), an elonging BC binding motif (*19*), and a cullin binding motif (*20*).

The TSC22D family is encoded in mammals by four different genes (TSC22D1-4), some of which can produce multiple isoforms. Of note, the TSC22D isoforms can be classified into short and long isoforms depending on the length of their N-terminal domain (Fig. 1A). Long isoforms contain a long and mainly disordered N-terminal domain (*11*, *12*), followed by a short, structured C-terminal domain known as TSC22 domain that is implicated in homo- and heterodimerization (*12*). Within the disordered N-terminal domain, a conserved region that mediates interaction with NRBP1 has been identified (*11*, *13*). Interestingly, within this region, in addition to a canonical RFxV motif, a highly conserved RWxC motif was also identified that participates in the interaction with NRBP1. Thus, it was proposed that CCT-binding motifs could be termed Rϕ (where ϕ represents an hydrophobic residue) (*12*). Short TSC22D isoforms lack most of the disordered N-terminal region, and thus lack Rϕ motifs, but retain the C-terminal TSC22 domain. Previous work has shown that short isoforms exert opposing effects to long isoforms (*13*).

**Figure 1.**
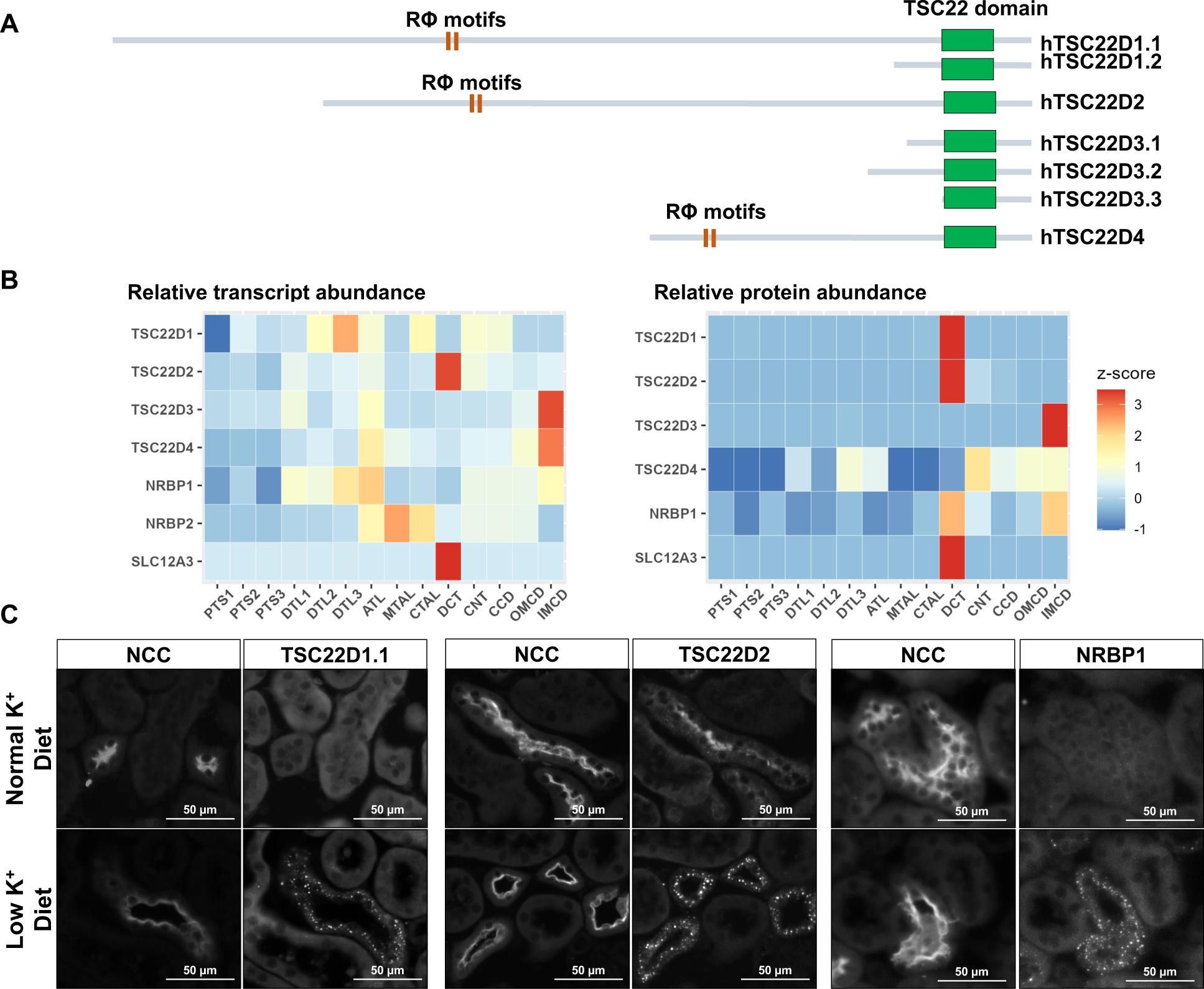
Within the kidney, TSC22D2 and TSC22D1.1 are enriched in DCT cells, and these proteins, as well as NRBP1, are observed in cytoplasmic puncta in DCTs of hypokalemic mice. **(A)** The TSC22D protein family. The four TSC22D genes (TSC22D1-4) present in mammals can give rise to multiple isoforms through alternative slicing. TSC22D isoforms can be classified into short and long, depending on the absence or presence, respectively, of a long, disordered N-terminal domain. The highly conserved C-terminal domain (named TSC22 domain) is present in all isoforms and is the only region of the protein that presents structural organization. This TSC domain is involved in protein homo- and heterodimerization (*12*). Interestingly, only the long isoforms present Rϕ motifs located within a region of intrinsic disorder that have been shown to be necessary for interaction with the CCT domain of NRBP1, but may also mediate interaction with SPAK/OSR1 and WNKs (*11–13*). **(B)** Relative abundance of transcripts and proteins of the NRBP and TSC22D families along different renal segments. To the left, a heatmap showing the relative abundance of NRBP and TSC22D transcripts along the 14 mouse nephron segments is presented. Data from single cell RNA sequencing experiments performed with manually microdissected segments produced by Chen and coworkers was used (*25*). To the right, a heatmap showing the relative abundance of NRBP and TSC22D proteins along the different rat nephron segments is presented. Data from quantitative proteomic experiments performed with manually microdissected nephron segments was used (*36*). Z-score normalization was applied to the data to standardize the values across each feature. **(C)** Immunofluorescent staining of kidney tissue from C57Bl/6 wild type mice maintained on normal or low K^+^ diet with antibodies against TSC22D2, TSC22D1.1 (the long isoform), and NRBP1. Identification of DCT cells was performed by co-staining with an antibody against NCC. Staining of tissue form at least three male or female mice per condition was performed with similar results in all cases.

Hyperosmotic stress has been shown to induce the formation of WNK1-containing biomolecular condensates that are formed through liquid-liquid phase separation in response to molecular crowding (*21*). The long, intrinsically disordered C-terminal domain of WNKs has been shown to be essential for the formation of WNK condensates. Interestingly, it has been recently shown that TSC22D and NRBP proteins also localize to WNK1-containing condensates induced by hypertonic stress (*11*). Moreover, it has been shown that, like WNK1, NRBP1 and TSC22D proteins are essential for adequate cell volume regulation in response to osmotic stress (*11*). In the kidney, and specifically in DCT cells, WNK condensates are observed under certain conditions in which WNK kinase abundance increases (e.g. hypokalemia and FHHt) (*22–24*).

Data available in kidney proteomic and transcriptomic databases show TSC22D2 mRNA and protein abundances are considerably higher in DCT than in other nephron segments (Fig. 1B). The data also reveal that the long isoform of TSC22D1 (TSC22D1.1) is also highly enriched within the DCT (*25*). In addition, knockout of the *TSC22D3* gene in mice (which encodes several short isoforms) leads to NCC overactivation and an FHHt-like phenotype (*26*). TSC22D3, also known as Glucocorticoid Induced Leucine Zipper (GILZ), has been shown to reduce SPAK and NCC phosphorylation when overexpressed in cells. These observations suggest that proteins from the TSC22D family in conjunction with NRBP, may play a relevant role in WNK signalling and DCT physiology. In the present study we explored the localization of specific TSC22Ds and NRBP1 in kidney tissues and investigated how these proteins impact the WNK4-SPAK/OSR1 pathway. Our data strongly suggest a physiological role of these proteins in DCT, and this role is most likely generalizable to other cell types and tissues where WNK function is essential.

## Results

### Expression of the long TSC22D isoforms TSC22D2 and TSC22D1.1 is enriched in the DCT where these proteins are present in WNK bodies together with NRBP1

A particular feature of the DCT is that, under certain conditions, WNK4, KS-WNK1, SPAK and OSR1 are observed to localize in cytoplasmic puncta (*22–24*) that are believed to comprise biomolecular condensates similar to the ones formed in cells treated with hypertonic conditions (*21*). These condensates have been termed WNK bodies. Thus, we sought to investigate the distribution of TSC22D2, TSC22D1.1, and NRBP1 proteins in kidney tissue by immunofluorescent staining at baseline and in mice maintained on a low K^+^ diet, a condition in which WNK bodies and NCC activation are observed (*22*, *24*). At baseline, no clear positive signal was observed with the TSC22D1.1 and NRBP1 antibodies, whereas the TSC22D2 antibody displayed a clear apical signal that was exclusively detected in NCC-positive cells (i.e. DCT cells) (Fig. 1C). In contrast, in kidneys from mice on low K^+^ diet, the three antibodies gave a similar signal: a cytoplasmic punctuate signal. This signal was mainly observed in DCT cells (Fig. 1C), but also seen in a few sporadic NCC-negative cells (Fig. S1). The latter were presumably CNT cells in which KS-WNK1 expression, a major driver of WNK body formation, is also observed (*22*, *27*). We cannot rule out that the absence of signal observed for NRBP1 and TSC22D1.1 in baseline conditions may have been due to insufficient sensitivity of the antibody assay, as NRBP1 expression, for example, is presumed to be ubiquitous and constitutive. However, detection within WNK bodies is likely to be made possible as a result of a high concentration of these proteins within these structures making these more readily detected in the immunofluorescence experiments. A punctuate cytoplasmic localization for TSC22D1.1 and TSC22D2 in DCT cells was also observed in other mouse models in which large WNK bodies have been previously reported: WNK4 knockout mice (*24*) and KLHL3-R528H knockin mice (*23*) (Fig. 2A).

**Figure 2.**
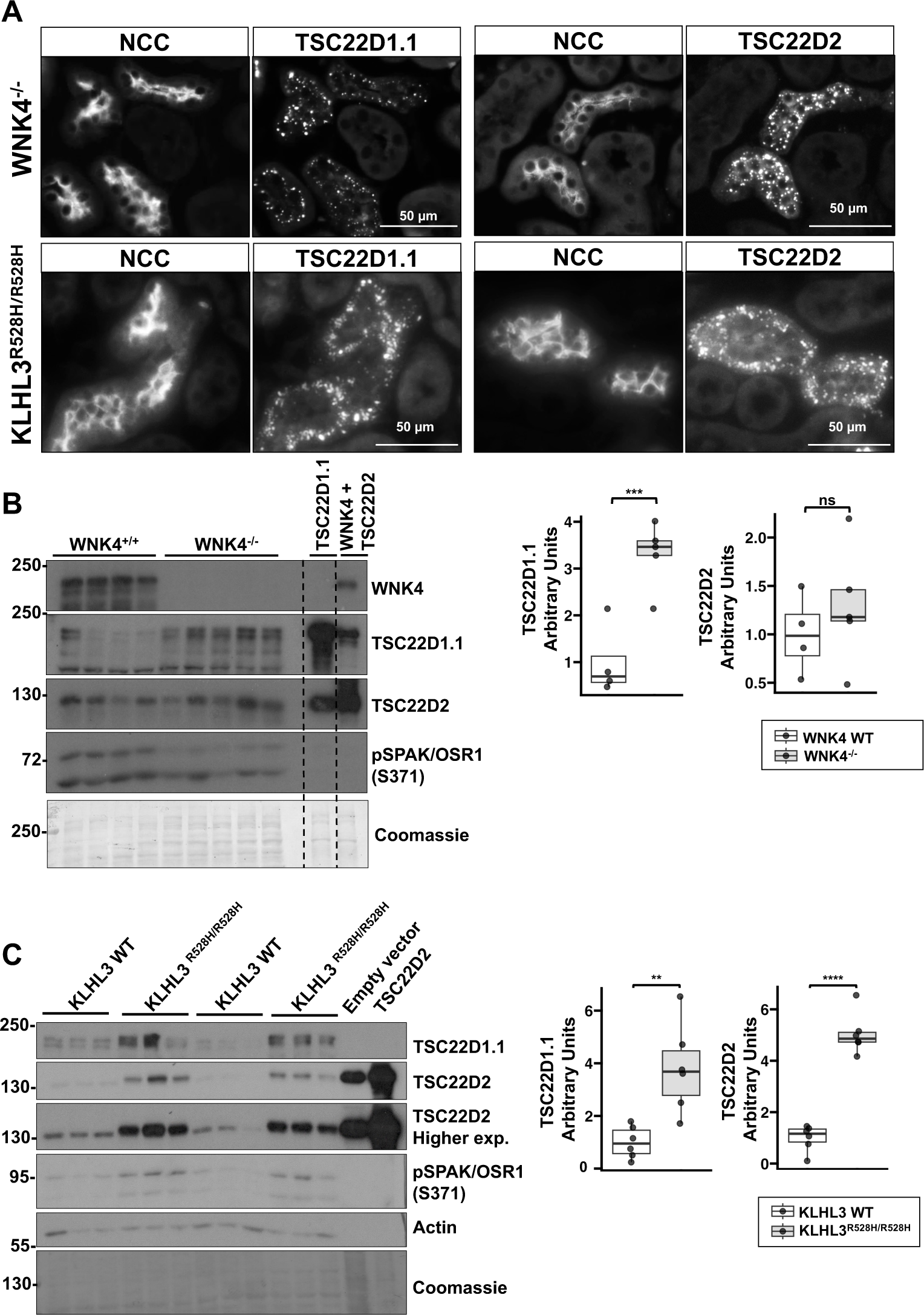
TSC22D2 and TSC22D1.1 are observed in cytoplasmic puncta in the DCTs of WNK4^-/-^ and KLHL3^R528H/R528H^ mice. **(A)** Immunofluorescent staining of kidney tissue from WNK4^-/-^ (top panels) or KLHL3^R528H/R528H^ (bottom panels) mice maintained on regular chow. TSC22D2 and TSC22D1.1-positive cytoplasmic puncta were observed almost exclusively in NCC-positive cells. Staining of tissue form at least three male or female mice per condition was performed with similar results in all cases. **(B)** Immunoblots performed with kidney samples from homozygous WNK4 knockout mice and their wild type littermates. The last two lanes were loaded with lysates from HEK293 cells transfected with TSC22D1.1 and TSC22D2 as control. Results of quantitation are presented in the graphs to the right. **(C)** Immunoblots performed with kidney samples from homozygous KLHL3-R528H mice and their wild type littermates. The last two lanes were loaded with lysates from HEK293 cells transfected with empty vector or TSC22D2 as control. Densitometric values were analyzed by unpaired Student’s t-test **p<0.01, ***p<0.001, ****p<0.0001.

Interestingly, greater TSC22D1.1 and TSC22D2 abundance was observed in KLHL3-R528H knockin mice and greater abundance of TSC22D1 was also observed in WNK4 knockout mice (Fig. 2B and C). In this latter model, WNK bodies presence is thought to be part of a failed compensatory response. Despite being described as transcriptional factors, we failed to detect nuclear signal of TSC22D1.1 or TSC22D2 in both transgenic models where the abundance of these proteins was increased.

We confirmed that TSC22D1.1, TSC22D2, and NRBP1-positive cytoplasmic puncta correspond to WNK bodies through colocalization with SPAK or WNK1, which was observed in tissues from mice on low K^+^ diet and from WNK4^-/-^ mice (Fig. 3). The signal observed with the WNK1 antibody most likely corresponded to KS-WNK1 (*23*, *28*). Altogether these results show that TSD22D1.1 and TSC22D2 within the kidney are mainly expressed in the DCT where these proteins colocalize with elements of the WNK-SPAK/OSR1 pathway in conditions in which WNK bodies are formed. In addition, NRBP1 is also present in DCT WNK bodies.

**Figure 3.**
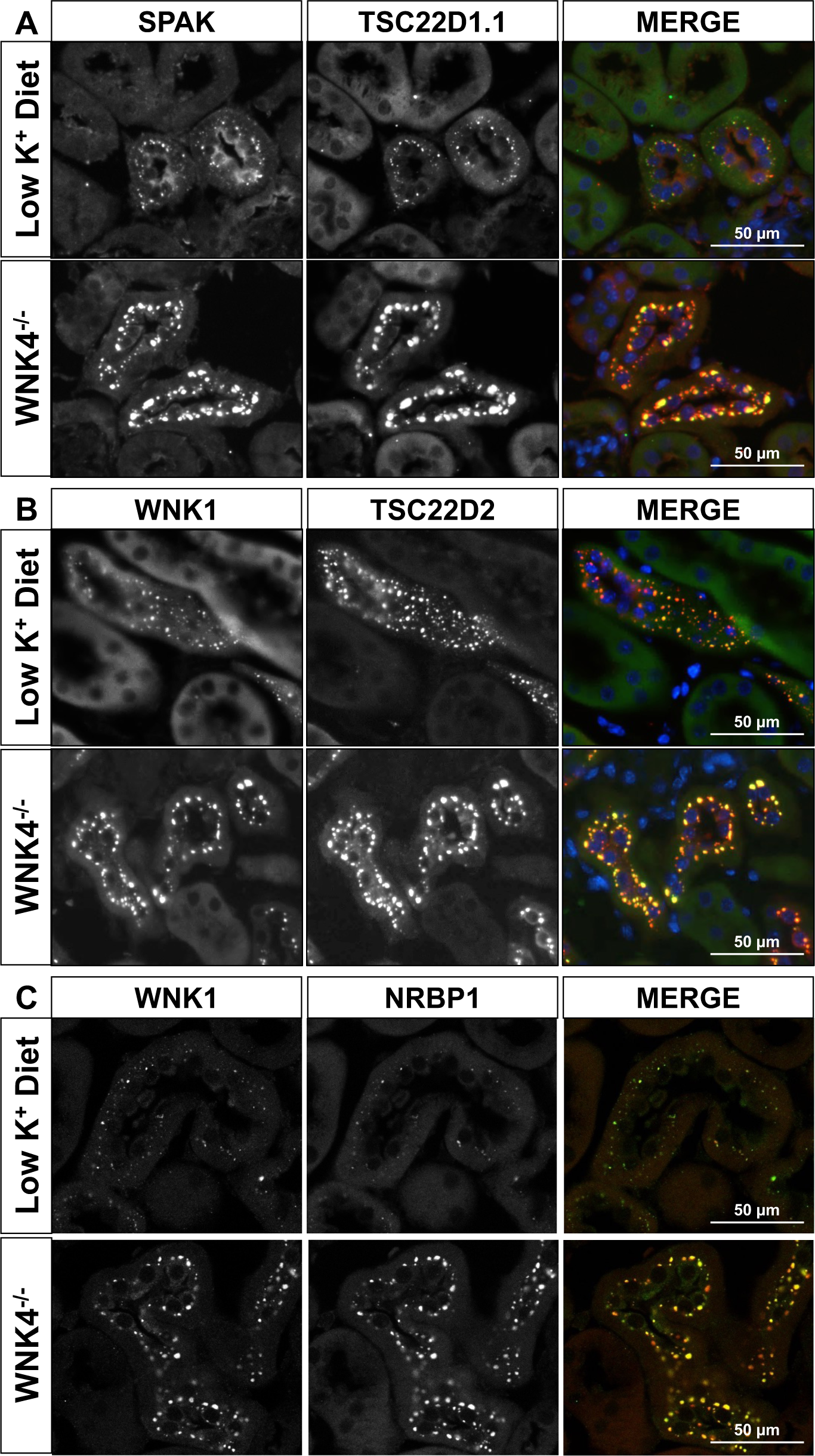
TSC22D2, TSC22D1.1, and NRBP1-positive cytoplasmic puncta are WNK bodies. Immunofluorescent staining of kidney tissue from C57Bl/6 wild type mice maintained on low K^+^ diet and WNK4^-/-^ on regular chow. Co-staining of TSC22D1.1 with SPAK **(A)**, TSC22D2 with WNK1 **(B)**, or NRBP1 with WNK1 **(C)** confirmed that TSC22D2, TSC22D1.1, and NRBP1-positive cytoplasmic puncta observed in DCT cells correspond to WNK bodies.

### Co-expression of NRBP1 with long TSC22D isoforms promotes activation of the WNK4-SPAK pathway

To explore the physiological role of TSC22D and NRBP proteins in the DCT, we decided to study the effect of these proteins on the activity of the WNK4-SPAK pathway. We focused on WNK4 because, as mentioned earlier, this is the main (and probably the only) catalytically active WNK kinase present in the DCT. HEK293 cells were co-transfected with SPAK, WNK4, NRBP1, and long TSC22D isoforms in different combinations (Fig. 4A and S2). Levels of SPAK phosphorylation (pSPAK) were assessed by immunoblot as a readout for WNK pathway activity. In the absence of transfected WNK4, TSC22D1 and TSC22D4, when expressed together with NRBP1, increased the levels of pSPAK (Fig. S2). A trend toward increased pSPAK was also observed in the presence of TSC22D2 and NRBP1. In the presence of WNK4, all three long TSCs, TSC22D1.1, TSC22D2 and TSC22D4, had an activating effect on SPAK phosphorylation when expressed in conjunction with NRBP1 (Fig. 4A and S2). Of note, activation of WNK4-SPAK was also observed in the presence of NRBP2/TSC22D2 (Fig.S3).

**Figure 4.**
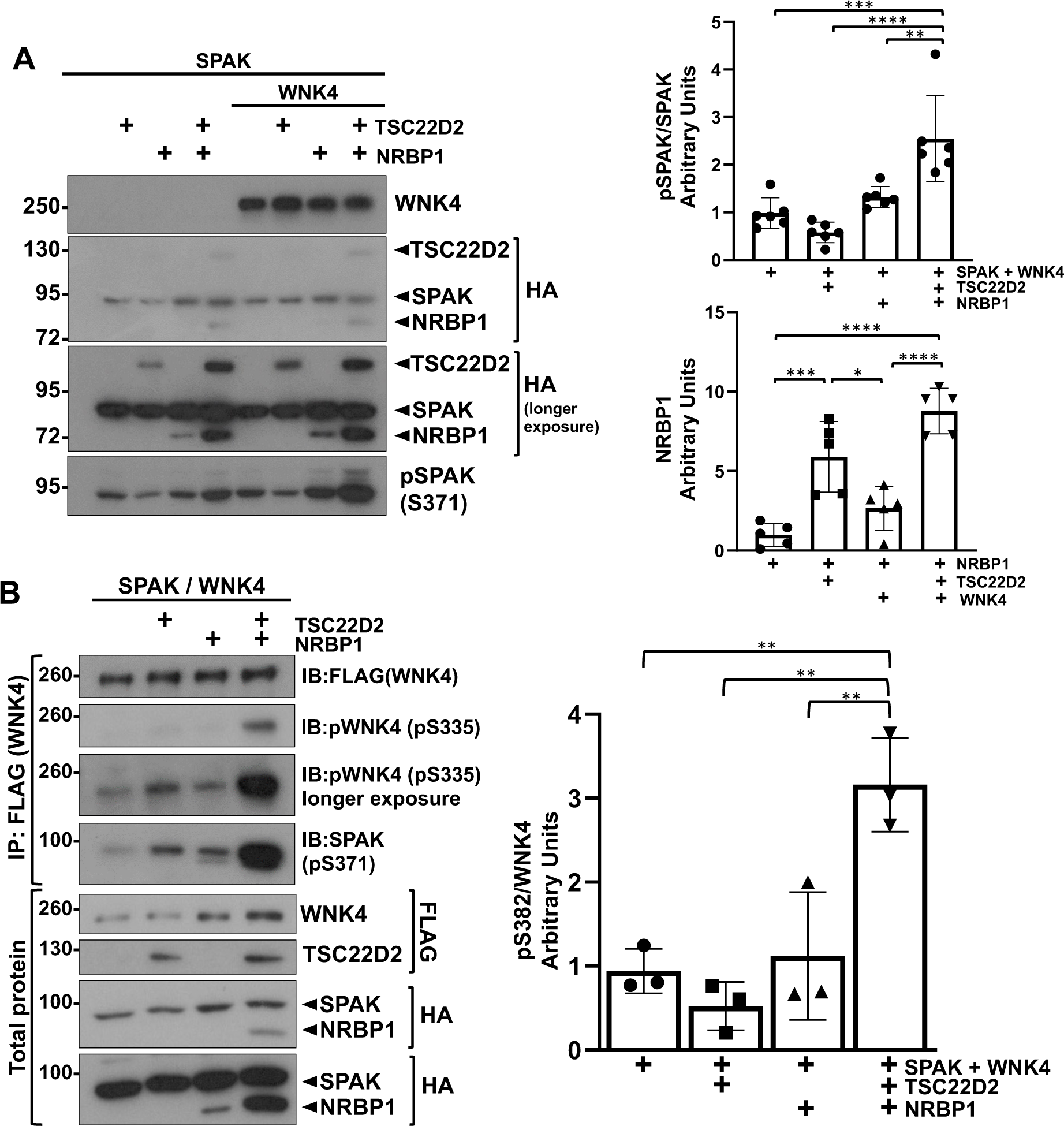
NRBP1 acts in concert with TSC22D2 to promote WNK4 activation. **(A)** The effect of co-expression of NRBP1 with TSC22D2 on WNK4’s ability to phosphorylate SPAK was assessed in HEK293 cells. Cells were transiently transfected with SPAK, WNK4, NRBP1 and TSC22D as indicated. 48 hours post transfection immunoblots were performed to confirm expression of transfected proteins and to analyze phosphorylation levels of SPAK (pSPAK-S371, which corresponds to the S motif site, previously identified as S373; see Q9UEW8 in Uniprot). An increase in pSPAK was observed upon co-expression of NRBP1 and TSC22D2 in the presence of WNK4. Results of quantitation are shown in the graphs to the right. ANOVA followed by Tukey post hoc tests were performed to identify statistically significant differences. *p<0.05, **p<0.01, ***p<0.001, ****p<0.0001. At least three independent experiments were performed. **(B)** HEK293 cells were transiently transfected with WNK4 and SPAK, as well as TSC22D2, NRBP1 or both. 48 hours post transfection autophosphorylation levels of WNK4 at the S335 activation site were assessed through immunoblot. A robust increase in WNK4-S335 phosphorylation was observed in cells co-expressing NRBP1 and TSC22D2. Results of quantitation are shown in the graph to the right. ANOVA followed by Tukey post hoc tests were performed to identify statistically significant differences. *p<0.05, **p<0.01, ***p<0.001, ****p<0.0001. At least three independent experiments were performed.

In these experiments, we noted that expression of long TSC22D proteins or WNK4 increased the amount of NRBP1 protein. This effect was additive, as higher NRBP1 levels were observed in the presence of a TSC22D and WNK4, than in the presence of either one of these proteins alone. The increase in pSPAK levels in the TSC22+NRBP1 groups was not due to the higher abundance of NRBP1 in this group because when we compared the effect of increasing NRBP1 abundance by increasing the amount of transfected DNA with the effect observed upon co-expression of TSC22D2, a much more notable increase in pSPAK was observed in the latter case (Fig. S4).

Finally, to investigate if the observed increase in pSPAK levels was due to increased activity of the WNK kinase (either the endogenous WNK or the transfected WNK4), we assessed WNK4 phosphorylation at the T-loop site (S335 in human WNK4; this is a site that is autophosphorylated and whose phosphorylation promotes kinase activation (*29*)). A clear increase in WNK4 T-loop phosphorylation was observed in the presence of TSC22D2 and NRBP1 (Fig. 4B), suggesting that these proteins increase the ability of WNKs to autophosphorylate. These findings are consistent with the observations made by us in a parallel work in which NRBP1 was shown to increase activity of WNK4 in in vitro kinase assays (*12*).

### Knockout of NRBP1 in the DCT reduces NCC phosphorylation

To confirm the role of NRBP1/TSC22Ds in the modulation of the WNK4-SPAK pathway in vivo, we generated DCT-specific, inducible NRBP1 knockout mice by crossing conditional ready NRBP1 mice (EMMA Strain ID: 09610) with NCC-Cre mice (*30*). Mice were studied on a normal K^+^ diet and on a low K^+^ diet for 5 days, the latter to promote WNK4-SPAK pathway activation in the DCT and the formation of WNK bodies. NRBP1 immunostaining of tissues from mice under low K^+^ diet confirmed the absence of NRBP1 expression in DCTs from tamoxifen-treated mice (Fig. 5A). NRBP1-positive condensates were observed in sporadic cells that were NCC-negative, supporting the specificity of the cell type-specific targeting strategy (Fig. S5). In contrast, NRBP2 staining in knockout mice on low K^+^ diet showed a clear positive signal in the DCT, where NRBP2 localized in WNK bodies. Large WNK4-positive WNK bodies were also observed in the knockouts. Moreover, under normal K^+^ diet WNK bodies were observed in the knockout mice, whereas, under these conditions, WNK bodies were absent in the control mice (Fig. 5B). Analysis of WNK4-positive WNK bodies of mice under low K^+^ diet showed that these were larger in size and had reduced circularity in the knockout mice (Fig. 5C). A tendency for higher number of WNK bodies was also observed. Interestingly, reduced pNCC and NCC levels were observed in NRBP1 knockouts (Fig. 5D-E). Previous work has also shown in WNK4 and other WNK-pathway deficient mice that loss of NCC phosphorylation results in a marked reduction in NCC protein (*7*, *31*). No differences in the levels of pSPAK/OSR1, SPAK, NRBP1, or WNK4 were observed presumably due the ubiquitous nature of these proteins that may have obscured changes occurring only in the DCT. Thus, the presence of NRBP2 in the DCT and the likely compensatory increase in WNK bodies observed in the knockouts did not appear to be sufficient to prevent a defect in pathway activation in the absence of NRBP1. This compensation, however, probably prevented a more dramatic decrease in NCC phosphorylation and activity which is why no dramatic electrolytic derangements were observed other than a tendency to hypokalemia (Table 1).

**Figure 5.**
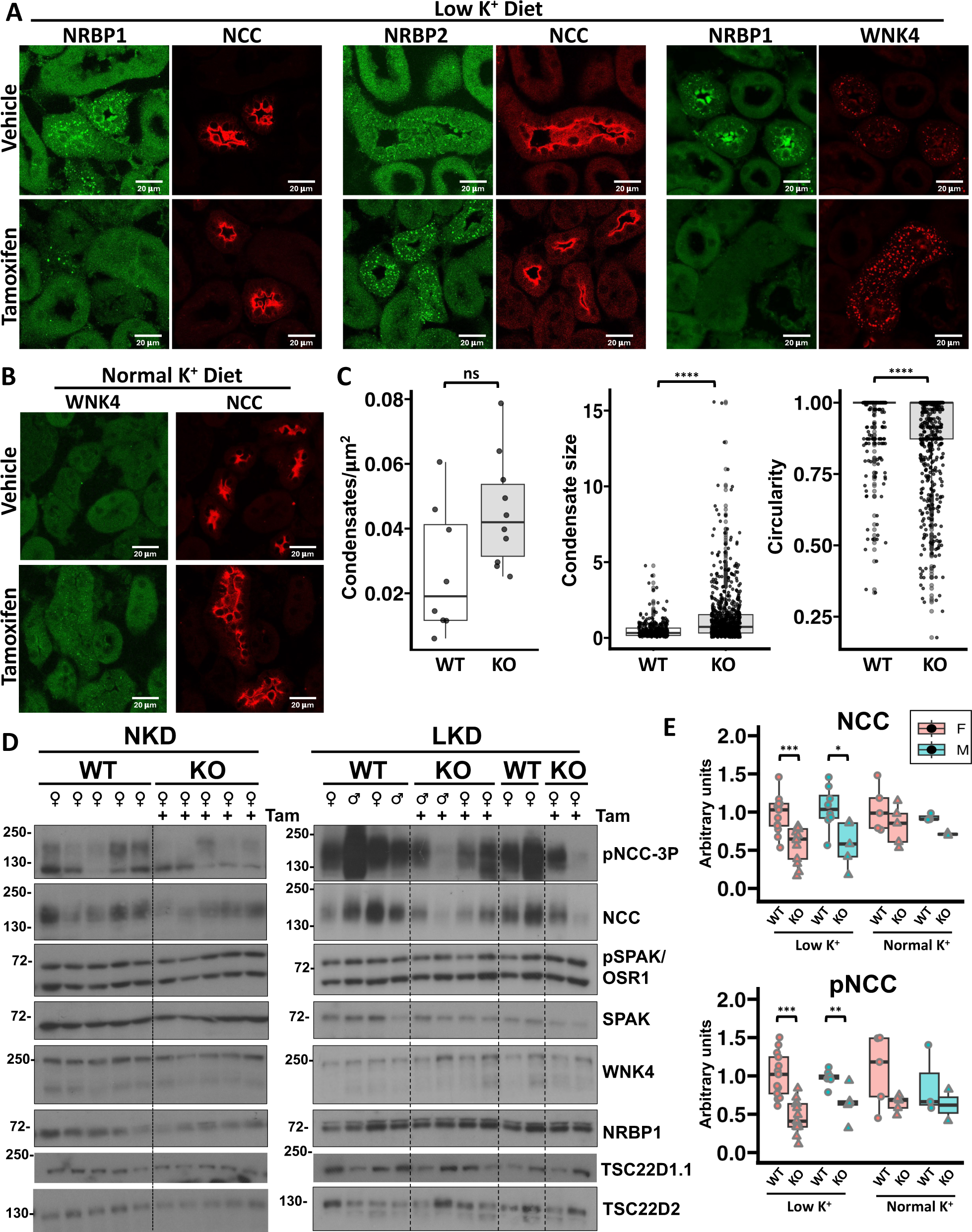
DCT-specific NRBP knockout mice have decreased NCC phosphorylation. **(A)** Deletion of NRBP1 in the DCT was confirmed by immunofluorescent staining of kidney sections from mice on low K^+^ diet (left panels). NCC positive tubules from knockout mice had no NRBP1-positive cytoplasmic condensates and the diffuse cytoplasmic signal was less intense than in other cell types. NRBP2 immunofluorescent staining revealed that NRBP2 is also present in DCT condensates from wild type mice on low K^+^ diet (middle panels). In knockout mice, NRBP2-positive condensates appeared to be larger. WNK4-positive cytoplasmic condensates in DCT cells from NRBP1 knockout mice also appeared to be larger. **(B)** Immunofluorescent staining of WNK4 and NCC in wild type and NRBP1 knockout mice maintained on normal K^+^ diet. No clear positive signal was observed for NRBP1 in wild type mice under this condition, whereas small WNK bodies were observed in DCTs of tamoxifen-treated knockout mice. **(C)** Analysis of number (left), size (middle), and circularity (left) of WNK4-positive WNK bodies observed in wild type and DCT-specific knockout mice was performed in Image J. ****p<0.0001 **(D)** Immunoblots performed with kidney protein lysates from wild type and DCT-specific NRBP1 knockout mice. Results of quantitation are of NCC and pNCC blots are presented in **(E)**. Student t tests were performed to identify statistically significant differences. *p<0.5, **p<0.01, ***p<0.001

**Table 1.** Plasma electrolytes of DCT-specific NRBP1 knockout mice and their wild type littermates when placed on a low K^+^ diet.

|  | Sex | Mean<br>NRBP1<sup>WT</sup><br>(mEq/L) | SD<br>NRBP1<sup>WT</sup><br>(mEq/L) | Mean<br>NRBP1<sup>KO</sup><br>(mEq/L) | SD<br>NRBP1<sup>KO</sup><br>(mEq/L) | n<br>NRBP1<sup>WT</sup> | n<br>NRBP1<sup>KO</sup> | p-<br>value | Significance |
| --- | --- | --- | --- | --- | --- | --- | --- | --- | --- |
| Na<sup>+</sup> | Female | 150.49 | 4.06 | 149.69 | 4.51 | 14 | 14 | 0.629 | ns |
|  | Male | 148.88 | 4.75 | 149.51 | 5.94 | 13 | 7 | 0.811 | ns |
| K<sup>+</sup> | Female | 2.68 | 0.6 | 2.37 | 0.47 | 14 | 14 | 0.141 | ns |
|  | Male | 2.69 | 0.6 | 2.17 | 0.5 | 13 | 7 | 0.055 | ns |
| Cl<sup>-</sup> | Female | 115.26 | 3.4 | 114.99 | 3.9 | 14 | 14 | 0.85 | ns |
|  | Male | 114.64 | 3.99 | 114.84 | 5.1 | 13 | 7 | 0.929 | ns |

### Activation of the WNK4-SPAK pathway by NRBP1/TSC22D2 depends on direct or indirect interactions mediated by CCT domains and Rχτ motifs

It is well known that the direct WNK-SPAK/OSR1 interaction depends on the CCT domain in SPAK/OSR1 and the Rϕ motifs present in WNKs (*15*, *32*, *33*). Interestingly, WNK kinases, in addition to Rϕ motifs, also contain two CCT-like (CCTL) domains termed CCTL1 and CCTL2 (Fig. 6A), whose function has remained elusive (*15*, *34*). The key residues within the CCT domain of OSR1 that establish interactions with residues of Rϕ motifs have been characterized (*17*). We have previously shown that mutations of some of these key residues in the CCTL domains of WNK4 (F476A, F478A) and (V701A, F703A) reduce the ability of the kinase to phosphorylate SPAK, without affecting its interaction with SPAK or its homo- or heterodimerization (*15*). Thus, we hypothesized that these CCTL domains of WNKs may be relevant for the interaction with the Rϕ motifs of long TSC22D proteins and that a reduced interaction with these proteins may explain the decreased activity of these mutants. To test this, we performed immunoprecipitation experiments in which we observed that, indeed, mutations within the first (CCTL1) or second (CCTL2) CCT motifs of WNK4 reduced its interaction with TSC22D2, and that mutations within both domains (CCTL1,2 mutant) further reduced the interaction (Fig. 6B).

**Figure 6.**
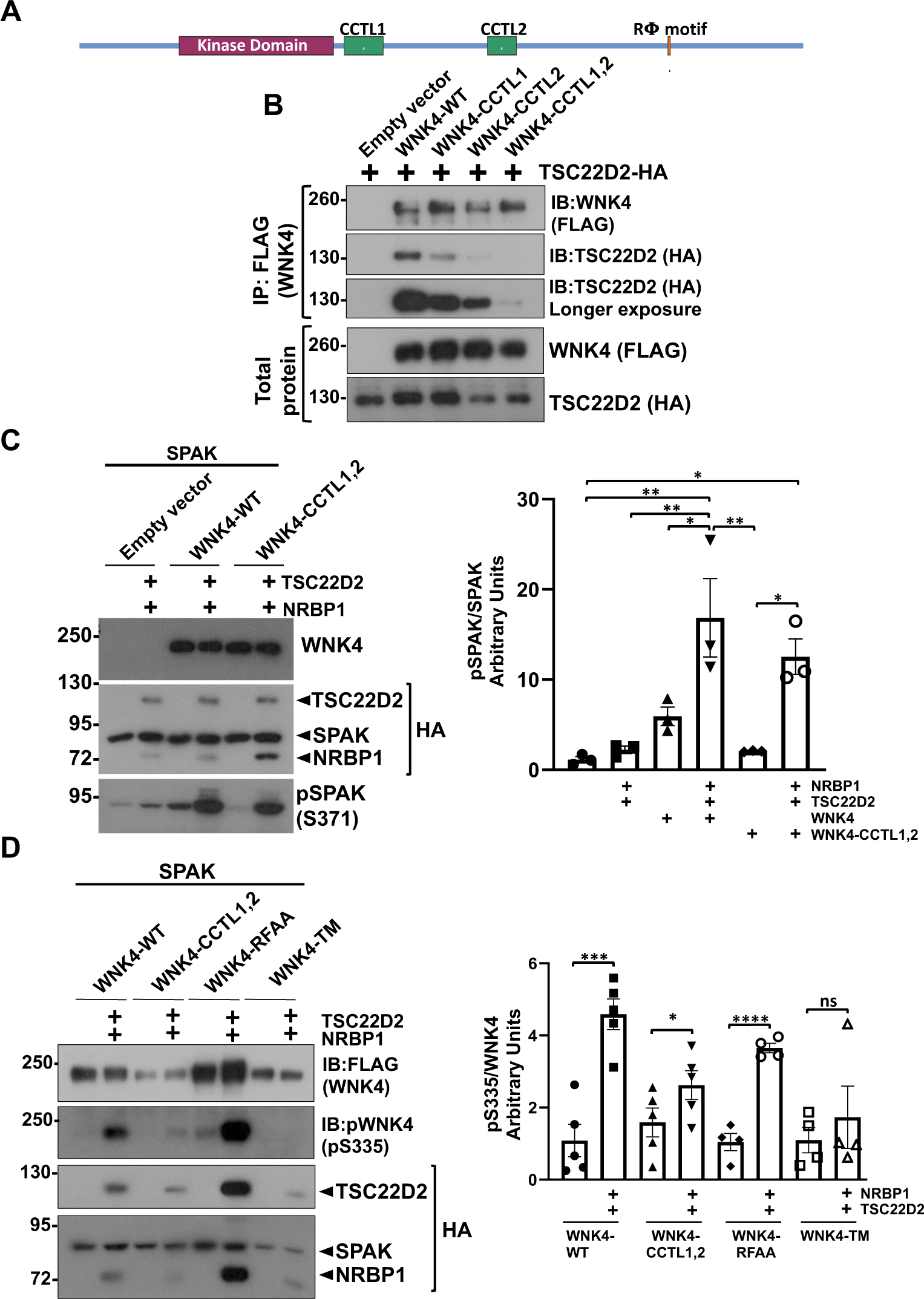
The activator effect of NRBP1/TSC22D2 on WNK4 depends on direct or indirect interactions between these proteins mediated by CCT domains and RFxV motifs. **(A)** Diagram depicting the primary structure of WNK4 in which the location of CCT domains and Rϕ motif are shown. Mutants harboring key mutations within these domains and motif were used for the experiments presented in this figure. **(B)** HEK293 cells were transiently transfected with TSC22D2 and wild type WNK4 or WNK4 mutants in which key residues within the first (CCTL1, F476A/F478A), second (CCTL2, F703A/F705A), or both (CCTL1,2) CCT domains present in WNK4 were mutated. 48 hours post transfection cells were lysed and TSC22D2 was immunoprecipitated. The levels of co-immunoprecipitated WNK4 were assessed by immunoblot. Significantly reduced WNK4-TSC22D2 binding was observed in the presence of mutations in both CCTL domains. **(C)** HEK293 cells were transiently transfected with SPAK and WNK4-WT or WNK4-CCTL1,2. In the indicated groups, NRBP1 and TSC22D2 were also co-transfected to assess their effect on WNK4-mediated SPAK phosphorylation. 48 hours post transfection pSPAK levels were analyzed by immunoblot. Despite the decreased TSC22D2-WNK4-CCTL1,2 binding observed in (B), activation of WNK4-mediated SPAK phosphorylation by NRBP1/TSC22D2 was observed in the presence of this mutant. **(D)** HEK293 cells were transiently transfected with SPAK and WNK4-WT, WNK4-CCTL1,2, WNK4-RFAA (with mutated RFXV motif necessary for direct SPAK binding (*15*), (R1016A, F1017A), or the triple mutant (WNK4-TM) containing the CCTL1,2 mutations and the RFAA mutations. In the indicated groups, NRBP1 and TSC22D2 were also co-transfected to assess their effect on WNK4 T-loop autophosphorylation. No increase in WNK4-S335 phosphorylation levels were observed upon NRBP1/TSC22D2 co-expression in the presence of the triple mutant that is defective in TSC22D2 binding and SPAK binding, suggesting that the activating effect observed in (C) in the presence of the WNK4-CCTL1,2 mutant was indirectly facilitated by SPAK. Results of blots quantitation are presented in the graphs to the right. ANOVA followed by Tukey post hoc tests were performed to identify statistically significant differences. *p<0.05, **p<0.01, ***p<0.001, ****p<0.0001. At least three independent experiments were performed.

To determine if the reduced activity of the WNK4 CCTL1,2 mutant was due to reduced interaction with TSC22D proteins, we examined the effect of NRBP1 and TSC22D2 co-expression on SPAK phosphorylation in the presence of this mutant version of WNK4. Unexpectedly, co-expression with NRBP1 and TSC22D2 significantly enhanced the mutant’s ability to phosphorylate SPAK (Fig. 6B). This led us to hypothesize that NRBP1 and TSC22D2 might activate WNK4-CCTL1,2 through an indirect interaction mediated by SPAK (i.e. TSC22D2 may recruit WNK4 through binding to SPAK). This was suspected given that absence of co-localization of TSC22D2 with the CCTL1,2 mutant was rescued in the presence of SPAK (Fig. S6). To test this hypothesis, we assessed WNK4 activation by measuring WNK4-T-loop phosphorylation in the wild-type protein, the WNK4-CCTL1,2 mutant, the WNK4-RFAA mutant (in which SPAK interaction is abrogated by mutating the Rϕ motif that mediates interaction with SPAK), and a triple mutant called WNK4-TM, in which both CCTL domains and the SPAK-binding site were mutated. This latter mutant was thus completely unable to establish CCT-Rϕ interactions. We observed that NRBP1 and TSC22D2 increased the level of T-loop phosphorylation for WNK4-WT, WNK4-RFAA, and WNK4-CCTL1,2, but this was not observed with the WNK4-TM construct (Fig. 6C). This suggests that the activation of the WNK4-CCTL1,2 mutant was indeed facilitated through an indirect interaction with NRBP1 and TSC22D2 mediated by SPAK. In summary, it would appear that there is a very large combinatory of Rϕ-CCT interactions in this pathway’s components that could partially compensate for the absence of a CCT domain or Rϕ motif in any single protein.

### Both, WNK proteins and long TSC22D proteins can promote condensate formation

When we observed subcellular localization of a GFP-tagged SPAK we noticed a diffuse cytoplasmic localization in the absence of WNK4 and TSC22D2. However, in the presence of WNK4 or TSC22D2, SPAK-GFP localized to cytoplasmic condensates. These condensates became more prominent when TSC22D2 and WNK4 were co-expressed (Fig. 7A). This suggests that WNK4 and TSC22D2 overexpression can promote condensate formation which is consistent with the fact that these proteins contain a large intrinsically disordered domains (*11*, *12*). Interestingly, no condensates were observed when SPAK, WNK4, and TSC22D2 were co-expressed with NRBP1 (Fig. 7B). Given that increased pSPAK levels are observed under this condition (Fig. 4) and given that the cells transfected with the four constructs had a more turgent appearance than cells transfected without NRBP1 (Fig. 7B), we suspected that absence of condensation could be due to an Slc12-mediated increase in cell volume that would decrease molecular crowding and prevent WNK condensation. This hypothesis was indeed supported by the observation that in Slc12 knockout cells, in which NKCC1 and all KCCs were knocked out and absence of SLC12 activity was confirmed (Fig. S7), NRBP1 overexpression in the presence of SPAK, WNK4, and TSC22D2 did not prevent condensate formation (Fig. 7B).

**Figure 7.**
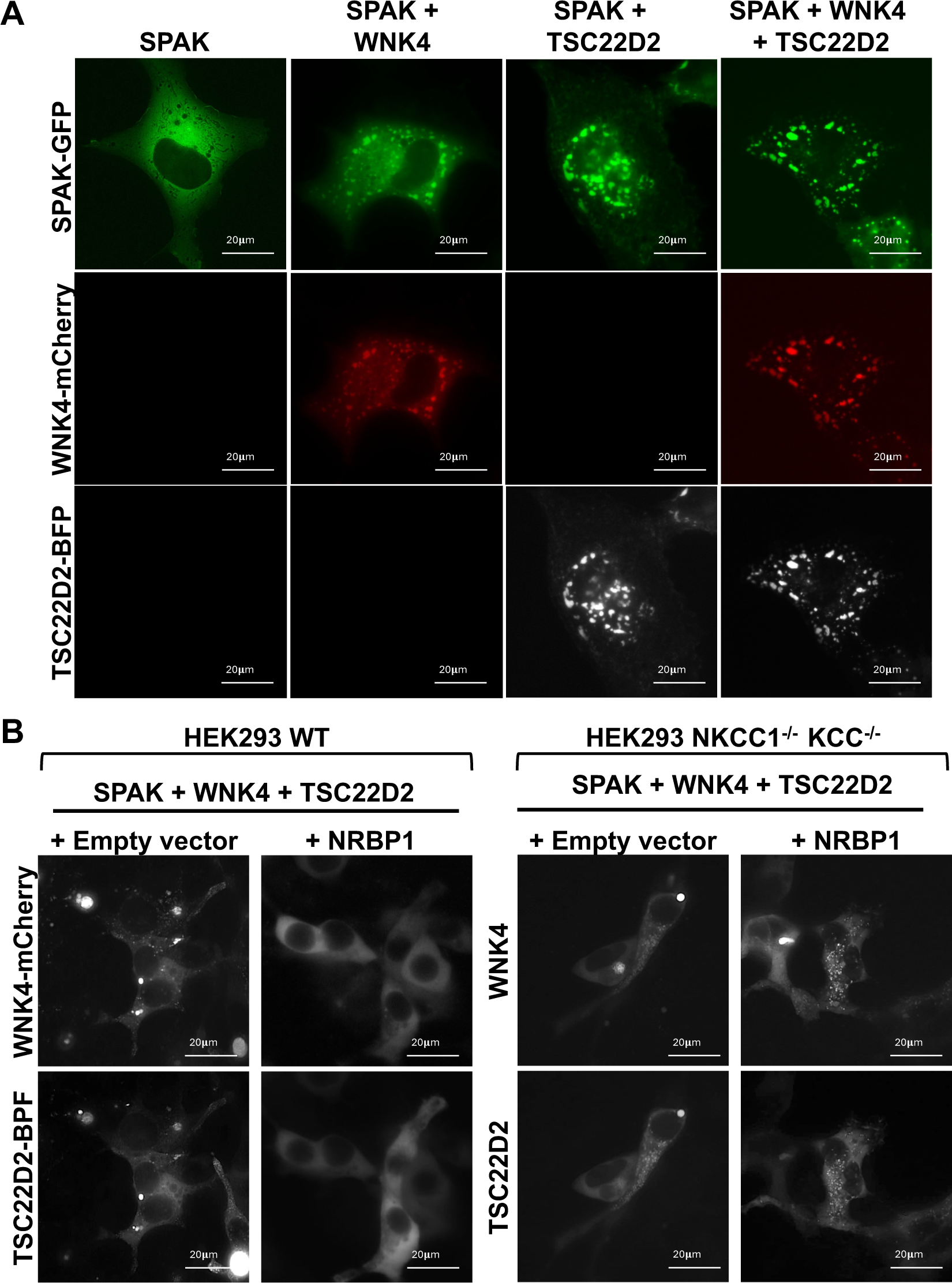
WNK4 and TSC22D2 expression in cultured cells promotes condensate formation and NRBP1 co-expression prevents condensate formation through an effect on cell volume. **(A)** COS7 cells were transfected with GFP-tagged SPAK and the indicated constructs. When SPAK-GFP was transfected alone, a diffuse cytoplasmic distribution was observed. In contrast, SPAK-containing cytoplasmic condensates were observed in the presence of overexpressed WNK4 or TSC22D2, and larger condensates were observed in the presence of both WNK4 and TSC22D2. **(B)** When WNK4-mCherry, SPAK-GFP, and TSC22D2-BFP were transfected in wildtype HEK293 cells, WNK4/TSC22D2—positive condensates were observed. However, when NRBP1 was also transfected, no condensates were observed and cells had a more turgent appearance. In contrast, in Slc12 knockout cells (see figure S7), condensates were observed in cells overexpressing the four proteins (i.e. including NRBP1), suggesting that the prevention of condensate formation observed in the presence of NRBP1 in HEK293 wild type cells was secondary to an effect on transport and most likely cell volume.

### Colocalization of WNKs and TSC22D proteins in condensates depend on both direct and indirect interactions

As mentioned above, the WNK4 T-loop phosphorylation experiments shown in Fig. 6C suggested the occurrence of an indirect interaction between WNK4 and TSC22D2 mediated by SPAK. Further supporting this observation, we observed co-localization of WNK4-WT, WNK4-CCTL1,2, and WNK4-RFAA with TSC22D2 and SPAK (Fig. 8A-C), but we did not observe colocalization of WNK4-TM with these proteins (Fig. 8D-E). Interestingly, in this latter case, we observed WNK4-positive condensates that were negative for TSC22D2 and SPAK, as well as TSC22D2/SPAK-positive condensates that were negative for WNK4 coexisting within cells (Fig. 8E). These results showed that, as long as WNK4 can establish interactions with TSC22D2 or SPAK, it can be recruited to TSC22D2 containing condensates (Fig. 8F). This also suggested that SPAK can maintain indirect interactions with WNK kinases through long TSC22D proteins. Altogether, these results suggest that any interaction that allows for recruitment of the necessary components to the condensates may be enough to facilitate the activation of this system. This conclusion was further supported by the following experiment. When the WNK4-TM was fused to FKBP12 and TSC22D2 was fused to the FKBP-Rapamycin Binding domain of mTOR (FRB domain), addition of rapamycin, a drug that induces the FKBP12-FRB interaction, induced co-localization of WNK4-TM and TSC22D2, as well as SPAK activation (Fig. 8G-J).

**Figure 8.**
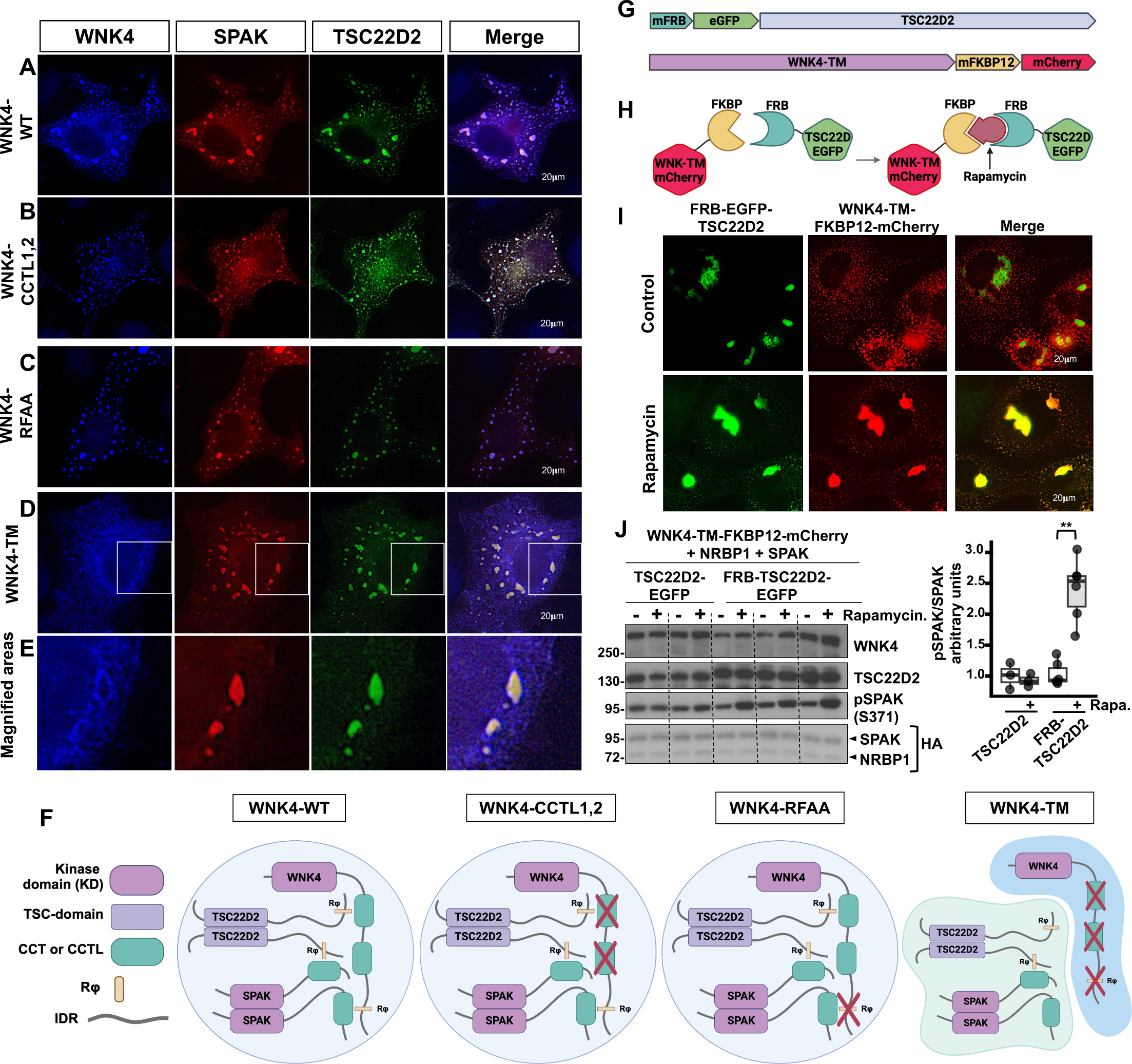
Colocalization of TSC22D2, WNK4, and SPAK within biomolecular condensates promotes pathway activation. COS7 cells were transiently transfected with SPAK-mCherry, TSC22D2-GFP, and wild type WNK4 **(A)** or the indicated WNK4 mutants **(B-D)**. Colocalization of WNK4 with TSC22D2 and SPAK in cytoplasmic condensates was observed with the wild type WNK4 (A) and was not affected by mutation of both CCTL domains in WNK4 (B) or by mutation of the Rϕ motif of WNK4 (C). However, when the three regions were mutated (CCTL1, CCTL2, and Rϕ), colocalization of WNK4 with SPAK and TSC22D2 was no longer observed (D-E). WNK4-containing condensates were observed, as well as SPAK-TSC22D2 containing condensates within a cell, that were mutually exclusive (D-E). (E) Magnification of the areas delimited with white boxes in figures shown in (E). Scale bars represent 20 um. **(F)** Cartoon that summarizes the observations made in experiments presented in (A-E). Wild type WNK4 can establish interactions with SPAK and TSC22D2 through its Rϕ motif and CCTL domains, respectively. When the CCTL domains of WNK4 are mutated (CCTL1,2), TSC22D can be recruited to WNK4-containing condensates via SPAK. When the Rϕ motif of WNK4 is mutated (WNK4-RFAA), SPAK can be recruited to WNK4-condensates via interaction with TSC22D2. However, when the CCTL domains and the Rϕ motif of WNK4 are all mutated (WNK4-TM), neither SPAK, nor TSC22D2 can be recruited to WNK4 condensates and independent TSC22D-SPAK containing condensates are formed. These observations are in accordance with the observation that the WNK4-TM construct cannot be activated by NRBP1/TSC222D2 (Fig. 6D). **(G)** Cartoon showing the mFRB-eGFP-TSC22D2 and WNK4-TM-mFKBP-mCherry constructs that were generated for the experiments presented in (H-I). **(H)** The WNK4-TM-mFKBP-mCherry protein is not expected to interact with mFRB-eGFP-TSC22D2 unless rapamycin is present. **(I)** Subcellular distribution of mFRB-eGFP-TSC22D2 and WNK4-TM-mFKBP-mCherry in the absence and presence of rapamycin. **(J)** Levels of pSPAK in the presence or absence of rapamycin were assessed by immunoblot. Results of quantitation are shown to the right. Student t test was performed to identify statistically significant differences. **p<0.01, n=5 from three independent experiments. Part of this figure was created in BioRender. Castaneda bueno, M. (2024) https://BioRender.com/e10z770 and https://BioRender.com/u96f731

### A short TSC22D isoform, TSC22D3.1, inhibits SPAK phosphorylation

Short TSC22D isoforms have been previously shown to exert opposite effects to TSC22D long isoforms (*13*). Moreover, knockout of *TSC22D3* in mice (which encodes for several short TSC22D isoforms) produces an FHHt-like phenotype with increased levels of phosphorylated NCC, thus suggesting that these short TSC22D isoforms may exert a negative effect on NCC (*26*). This motivated us to investigate the effect of TSC22D3 on the WNK4-SPAK pathway. We observed that the expression of TSC22D3.1 decreased the phosphorylation of SPAK in cells expressing SPAK, WNK4, and NRBP1 (Fig. 9A-B). A similar inhibitory effect of TSC22D3.1 was observed in the presence of overexpressed TSC22D2.

**Figure 9.**
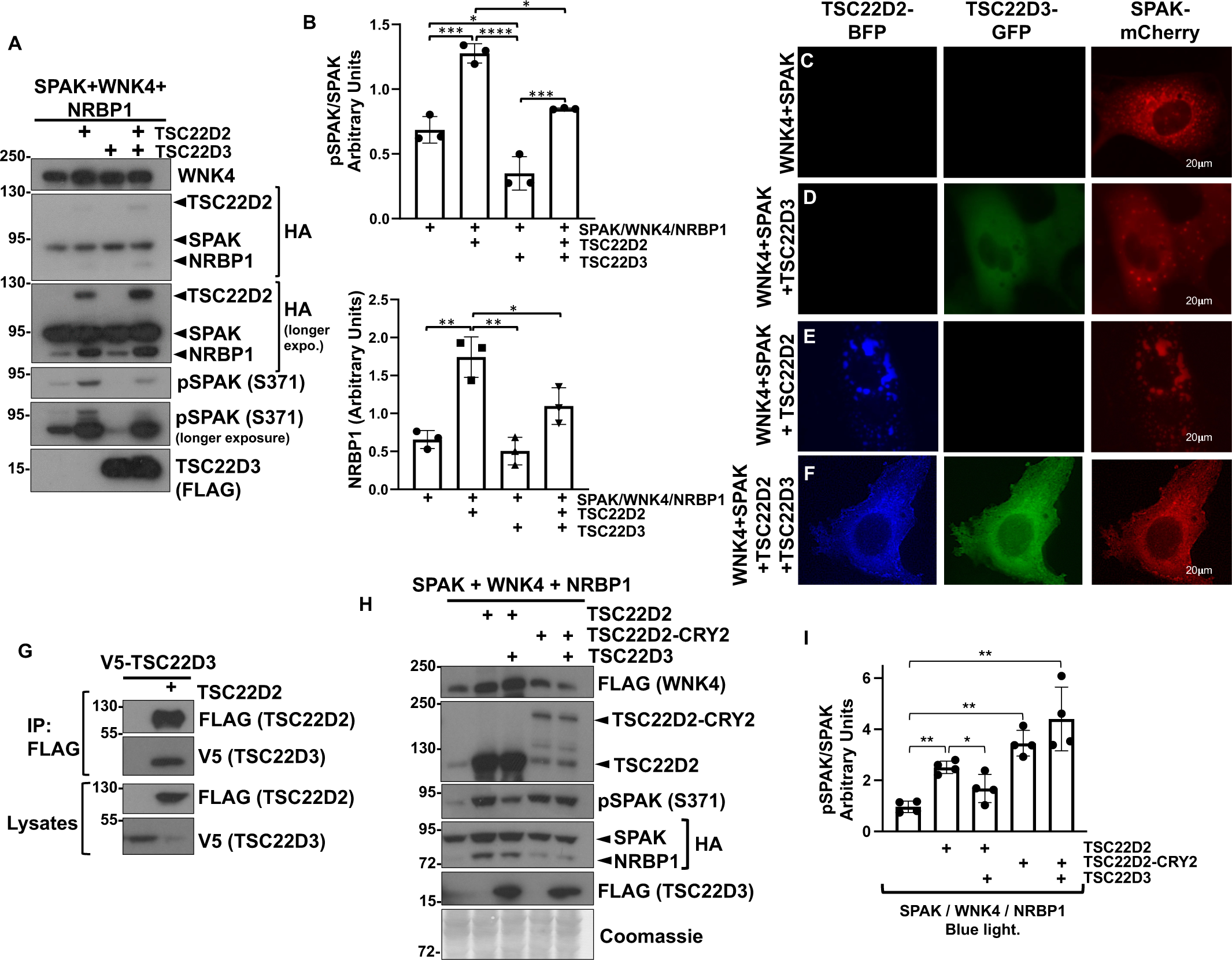
TSC22D3.1, which is a short TSC22D protein, exerts an inhibitory effect on the WNK-SPAK pathway. **(A)** HEK293 cells were transiently transfected with SPAK, WNK4, NRBP1, TSC22D2, or TSC22D3 as indicated. 48 hours post transfection pSPAK levels were analyzed by immunoblot. **(B)** Results of quantitation of blots represented in (A). ANOVA followed by Tukey post hoc tests were performed to identify statistically significant differences. *p<0.05, **p<0.01, ***p<0.001, ****p<0.0001. At least three independent experiments were performed. **(C-F)** Microscopic images showing cellular distribution of TSC22D2-BFP, TSC22D3-GFP, and SPAK-mCherry. Cells were transfected with the constructs indicated to the left. Co-expression of SPAK with TSC22D3, TSC22D2, or both affects the localization of SPAK within the cell. TSC22D2 induce the formation of larger, and more irregular condensates, whereas addition of TSC22D3 reduces the size of condensates and favors a diffuse cytoplasmic localization of SPAK. **(G)** Co-immunoprecipitation assays performed with lysates from HEK293 cells transfected with the indicated constructs. FLAG-tagged TSC22D2 was immunoprecipitated and binding of TSC22D3-V5 was assessed by immunoblot. **(H)** TSC22D2 self-oligomerization prevents the inhibitory effect of TSC22D3. A TSC22D2-Cry2 fusion construct was used for these experiments. The Cry2-Clust domain fused to TSC22D2 is known to oligomerize in response to stimulation with blue light. HEK293 cells were transfected with the indicated constructs. 48 hours after transfection cells were stimulated with blue light for 30 minutes and then lysed. SPAK phosphorylation levels were assessed by immunoblot. **(I)** Results of quantitation of blots presented in (H). ANOVA followed by Tukey post hoc tests were performed to identify statistically significant differences. *p<0.05, **p<0.01. At least three independent experiments were performed.

To assess the differential and/or possibly opposing effect of a short versus a long TSC22D protein on WNK condensate formation, we expressed TSC22D2 (a long TSC22D isoform) and TSC22D3.1 (a short TSC22D isoform) alone or in combination, together with WNK4 and SPAK, and observed their effects. When cotransfected with WNK4 alone, SPAK was localized in cytoplasmic condensates (Figure 7C). SPAK localization was similar when TSC22D3 was added, while TSC22D3 showed a diffuse cytoplasmic localization (i.e. not in WNK condensates) (Fig. 9D). In contrast, overexpression of TSC22D2 with SPAK and WNK4 (in the absence of TSC22D3) resulted in condensates in which SPAK and TSC22D2 colocalized. These were larger and more irregular in shape, supporting a pro-condensation role, or a condensate modifying property for long TSC22D proteins (Fig. 9E). Finally, in the presence of all components (WNK4, TSC22D3, and TSC22D2), SPAK localization was diffuse in most cells (Fig. 9F). However, condensates were still present in some cells, and these condensates were positive for both TSC22D2 and TSC22D3 (not shown).

We also observed that TSC22D3 can interact with TSC22D2 in immunoprecipitation assays (Figure 9G). This interaction is most likely mediated by the TSC22D heterodimerization domain as suggested by data from our parallel work ((*12*), Fig. 10) and thus we hypothesize that prevention of TSC22D2 dimerization in the presence of TSC22D3 may explain the inhibitory effect of the latter. This hypothesis was supported by the observation that stimulation of TSC22D2 self-oligomerization prevented the inhibitory effect of TSC22D3. Self-oligomerization of TSC22D2 was achieved by fusing TSC22D2 to Cry2-Clust, a light-responsive oligomerization domain from *A. Thaliana* (*35*) (Fig 9H).

**Figure 10.**
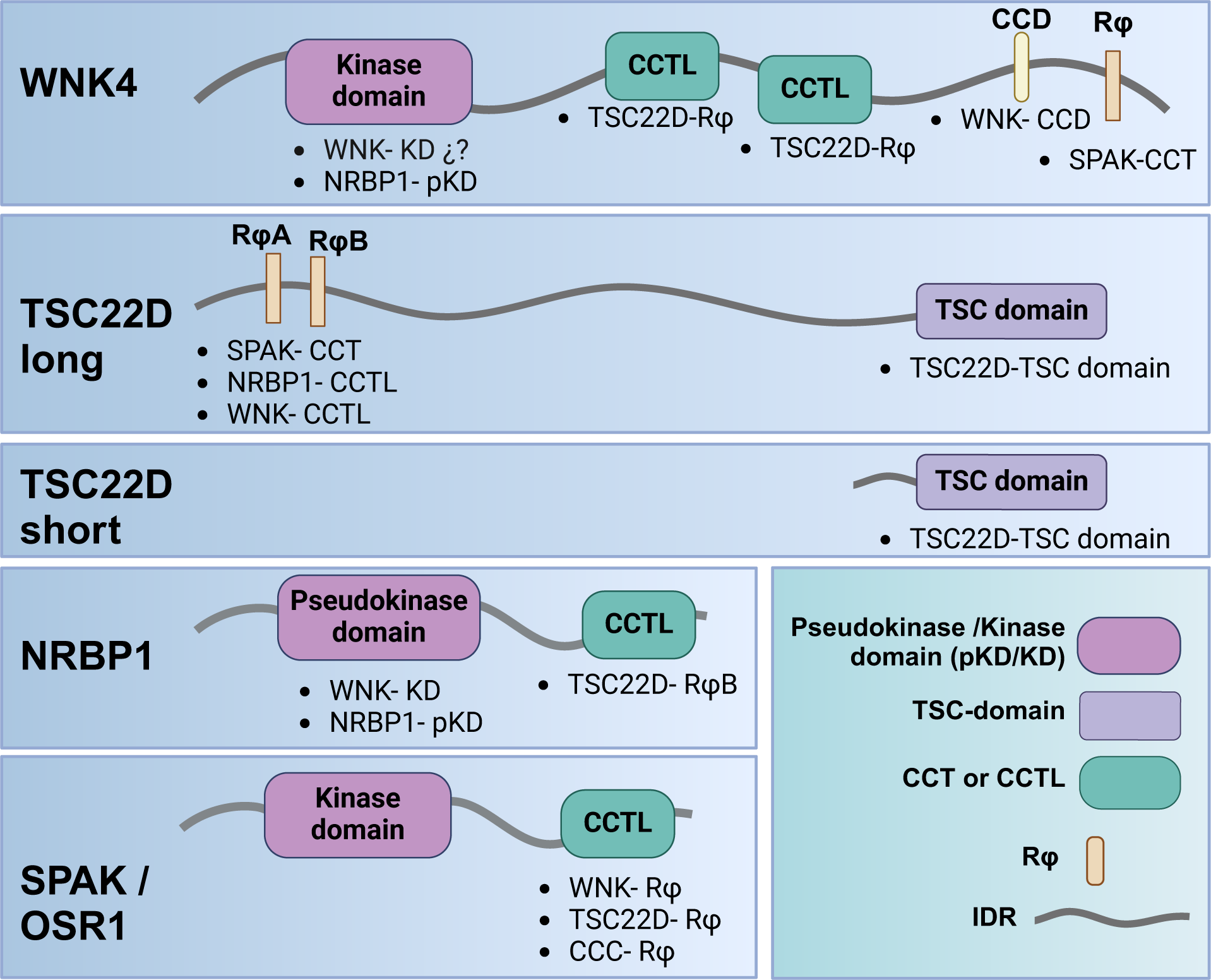
Motifs and domains mediating interactions between components of the WNK-SPAK/OSR1-TSC22D-NRBP pathway. Below the indicated domains and motifs, we list the specific domains or motifs with which there is experimental validation of interaction or prediction of interaction from structural models. **WNKs c**ontain one or more Rϕ motifs within their C-terminal domain that interact with the CCT domain of SPAK (*15*, *16*, *32*). WNKs also contain two CCTL domains in their C-terminus that mediate interaction with Rϕ motifs in long TSC22D proteins (this work and (*12*)). The absence of this interaction may explain the loss of function effect of mutations within these domains (*15*, *46*). Finally, WNKs can homo- and heterodimerize via their C-terminal coiled-coil domain (*47*) or through kinase domain-mediated interactions (*48*). WNKs can also interact with NRBP1 apparently via a kinase-pseudokinase domain mediated interaction (*12*). **SPAK** can also interact with CCCs through a SPAK CCT-domain -CCC-Rϕ interaction (*49*, *50*) and with TSC22Ds through a SPAK-CCT-domain - TSC22D-Rϕ interaction (*51*). We speculate that the presence of TSC22D2 (Fig. 1) and NRBP1 (Fig. 5) in the apical membrane of DCT cells may be due to this SPAK-TSC22D interaction. **Long TSC22D proteins** can also interact with the CCT domain of NRBP1 via an Rϕ motif (*11–13*). Finally, **long and short TSC22D proteins** can homo and hetero-dimerize via their C-terminal TSC domain (*12*). We speculate that the inhibitory effect of TSC22D3 on the WNK-SPAK/OSR1 pathway (Fig. 9, (*26*)) may be due to its heterodimerization with long TSC22D proteins, thus somehow preventing their activating effect. Created in BioRender. Castaneda bueno, M. (2024) https://BioRender.com/c08f700

In summary, our observations suggest that short TSC22D isoforms may oppose the effects of long TSC22D isoforms on the WNK-SPAK/OSR1 pathway. In our system, TSC22D3 appeared to oppose the pro-condensation effect of TSC22D2 (perhaps through heterodimerization), and this effect may be related to its negative effect on pathway activation.

## Discussion

The very recent identification by us and an independent group of the functional interaction of NRBP1 and TSC22D proteins with the WNK-SPAK-CCC pathway (*11*)(Alessi) motivated our study of the role of these proteins in the regulation of WNK signaling, NCC activity, and DCT physiology. First, we showed that TSC22D2 and TSC22D1.1 expression is indeed enriched in the DCT when compared to other nephron segments (Fig. 1), consistent with previous transcriptomic and proteomic studies. This further highlights the relevance of this signaling pathway in these cells where WNK4 and KS-WNK1 are also enriched (*25*, *36*). Moreover, our data showed that, under different conditions in which WNK bodies are formed in the DCT and NCC is activated (e.g. low K^+^ diet), TSC22D1.1, TSC22D2, and NRBP1 localize within WNK bodies. This constitutes an additional piece of evidence suggesting the similarity of the low K^+^-induced DCT WNK bodies and the WNK condensates that are induced by hypertonicity in different cell types as part of the cell-volume regulatory response (*11*).

Second, our data suggest that NRBP1 and TSC22D proteins can exert an activating effect on the WNK4-SPAK/OSR1 pathway by promoting an increase in the autophosphorylation of WNK4 (Fig. 4). This is consistent with the observation by Xiao et al. that, like WNK1, TSC22D2 and NRBP1 also participate in cell volume regulation and are functionally related as revealed by co-essentiality analysis (*11*). It is also consistent with the observation that NRBP1 knockout in cultured cells reduces pSPAK and phosphorylation of WNK1 at the activating Ser382 phosphorylation site, and also that, in in vitro kinase assays performed with recombinant WNK4 (1–449), NRBP1 presence increased the T-loop autophosphorylation of WNK4 as well as the WNK4-mediated OSR1 phosphorylation (*12*). This latter result, however, is slightly different from our observations made in the HEK293 assay because in these cells we only observed WNK4 activation in the presence of both NRBP1 and long TSC22D proteins, but not in the presence of NRBP1 alone (Fig. 4). This was particularly evident in the dose-response experiments in which the addition of increasing amounts of NRBP1 did not increase pSPAK levels, but addition of NRBP1 together with TSC22D2 had a robust activating effect (Fig. S4). The contrast in these observations is likely due to the different nature of the assays (e.g. in vitro context vs. cellular context). Our studies indicate that TSC22D isoforms are likely to be required for NRBP1 to maximally activate WNK4 and likely other WNK isoforms. In future work, it would be important for the impact of TSC22D isoforms to be assessed in the in vitro activation studies.

We speculate that a direct interaction between the WNK kinase domain and the NRBP1 pseudokinase domain may occur and that, like for other pseudokinases that have regulatory activity on their related kinases, the pseudokinase (NRBP1) may adopt an active conformation in response to a stimulus (e.g. hypertonicity) that permits its interaction with the kinase (WNKs) leading to its allosteric activation. This proposal is based on interaction models presented in a parallel study (*12*), but importantly this hypothesis remains to be tested experimentally. The requirement for both a long TSC22D protein and NRBP1 to observe activation in the cellular HEK293 assay suggests that TSC22D interaction with NRBP1 may be required to achieve the conformation necessary for interaction with WNK in vivo. Another possibility is that, given that TSC22D proteins are highly disordered and appear to contribute to condensate formation (Fig. 7), overexpression of both NRBP1 and long TSC22Ds promote the WNK-NRBP1 interaction to occur within biomolecular condensates where the high concentrations of components of this pathway may enhance WNK kinase activation. Alternatively, WNK condensates may constitute an environment with specific physicochemical properties that may favor WNK activation by NRBP1.

The modulatory role of NRBP1 and TSC22D proteins on the WNK-SPAK/OSR1 pathway was demonstrated in vivo, as the inducible deletion of NRBP1 specifically in the DCT significantly reduced the phosphorylation of NCC (Fig. 5). NRBP2 presence in these cells likely exerts some compensatory effect, as our data show that NRBP2 can also promote WNK-SPAK/OSR1 activation (Fig. S3), and NRBP2 presence was clearly observed in DCT cells (Fig. 5). Moreover, increased WNK bodies’ size in the knockout mice was probably also part of a compensatory response. Increased presence of WNK bodies has been observed in other models with reduced NCC activity like WNK4 knockout mice and CAB39/CAB39L double knockout mice (*24*, *37*). Interestingly, in the present study increased TSC22D1.1 kidney abundance was observed in WNK4 knockout mice which might also be part of the failed compensatory response that occurs in these mice.

Previous works have shown other functional roles of NRBP1 and TSC22D proteins. For example, NRBP1 has been proposed to function as a substrate adaptor molecule for Cullin-Ring Ligase (CRL) complexes with ubiquitin ligase activity (*20*, *38*). Whether this function is related to their ability to modulate the WNK-SPAK/OSR1 signaling pathway remains to be determined. However, the observation that NRBP1 binding to CRL elements (e.g. Elongins B and C) is not modulated by hypertonicity like interaction with WNKs (*12*), suggests that these may constitute independent functions. One of the proposed targets for a NRBP1-containig CRL complex is SALL4, a transcription factor that is key in mediating stem cell fate (*38*, *39*). Inducible knockout of NRBP1 in intestinal progenitor-like cells of mice caused aberrant proliferation of these cells and SALL4 was found to be increased (*38*). Interestingly, a SALL4 paralogue, SALL3 is highly enriched in the DCT (*25*, *40*). Our data shows that SALL3 can indeed bind NRBP1 in vitro (Fig. S8), but we did not observe changes in SALL3 expression or localization in the DCTs of NRBP1 knockout mice (Fig. S8). Further work is necessary to explore the possible role of NRBP1 in SALL3 regulation in the DCT. In addition, the evidence of the role of NRBP1/TSC22Ds in the regulation of cell proliferation is extensive. For instance, global knockout of NRBP1 in mice was observed to produce an increased incidence of tumorigenesis (*38*) and decreased NRBP1 expression has been observed in a wide variety of tumors (*38*, *41*). In drosophila, the NRBP1 homologue MADM and the long TSC22-like protein Bunched A have been shown to promote growth (cell number and cell size) (*13*, *42*). Whether these roles are related to WNK modulation, CRL function, or both remains uncertain. Further exploration of the DCT-specific knockout mice will be necessary to assess if NRBP1 absence affects DCT cell proliferation.

Finally, we also investigated the role of specific domains and motifs involved in the interactions among the components of this pathway. Mainly, we focused on defining the role of each interaction domain/motif present in WNK4. The presence of a single validated Rϕ motif in WNK4 simplified this analysis. Our data is consistent with the notion that the CCT domain-Rϕ motif interaction mechanism comprises the dominant mechanism mediating site-specific interactions among components of this pathway. A summary of the current knowledge is presented in Fig. 10. An Alphafold model has been generated in which most of these experimentally-validated interactions are predicted (*12*). It remains to be determined whether this structured complex can exist within WNK biomolecular condensates. Our data provides at least some evidence that there is certain flexibility regarding the specific interactions that can be used to recruit pathway components to the condensates and promote pathway activation. For instance, it was observed that, in the absence of direct WNK-TSC22D interaction (WNK4-CCT1,2 mutant), the SPAK-WNK4 interaction could recruit WNK4 into TSC22D2 containing condensates, and thus, TSC22D/NRBP1-mediated activation of WNK4 could occur (Figs. 6 and 8). Furthermore, fusion of the exogenous interaction modules FRB and FKBP12 to TSC22D2 and to the WNK4 triple mutant who is incapable of binding SPAK or TSC22Ds, respectively, rescued the interaction in the presence of rapamycin and promoted pathway activation (Fig. 8F-I).

## Materials and Methods

### Mice studies

Animal studies were approved by the Animal Care and Use Committee of the Instituto Nacional de Ciencias Médicas y Nutrición Salvador Zubirán. For immunolocalization studies, male and female wild type mice were given low K^+^ diet (LKD, 0% K^+^, Research Diets D16120202Mi) or normal K^+^ diet (NKD, 0.8% K^+^, that was prepared by adding KCl to the LKD) for 7 days. KLHL3-R528H homozygous mice (*23*) and WNK4-knockout homozygous mice (*7*) were also studied under baseline conditions (normal chow).

DCT-specific, inducible knockout mice were generated as follows. Sperm from the C57BL/6N-A<TM1BRD>Nrbp1<tm3a(EUCOMM)Wtsi>/WtsiOulu strain was obtained from the EMMA repository (EMMA Strain ID: 09610). IVF was performed to rederive the strain (fertilization and implantation of C57BL6/J oocytes into pseudopregnant C57BL/6J females). Heterozygous progeny carrying the 3 loxP Nrbp1 allele (identified by PCR of tail genomic DNA) were then crossed with FLP-e knockin mice (Jackson Laboratories, stock # 016226, B6N.129S4-Gt(ROSA)^26Sortm1(FLP1)Dym^/J) to eliminate the neomycin-cassette, the third loxP site, and generate the conditional ready allele (TM3c, https://www.eummcr.info/faq#nomenclature_eucomm). After confirming FlpE-mediated recombination, mice harbouring the conditional ready allele were bred with NCC-Cre mice (*30*). For experiments, homozygous NRBP1 floxed mice that were also heterozygous for the NCC-Cre allele (NRBP^fl/fl^;NCC^CreERT/-^) were treated with five doses of tamoxifen (100 mg/kg/d) given every other day to induce the deletion (*30*). Wild type controls were NRBP^wt/wt^;NCC^CreERT/-^, NRBP^wt/fl^;NCC^CreERT/-^, or NRBP^fl/fl^;NCC^CreERT/-^ that were treated with vehicle (100 μl / 20g of body weight of sterile corn oil (Thermo Sceintific, #AC405435000)). After tamoxifen treatment, low K^+^ diet (0% K^+^) was given to the mice for five days, after which mice were euthanized.

Tissue collection was performed in the following way. Animals were anesthetized with isoflurane (2%). The left kidney was harvested for immunoblot studies. The right kidney was perfused with 20 ml of PBS followed by 20 ml of 4% (w/v) paraformaldehyde in PBS for immunofluorescence studies. Harvested kidneys were incubated for 3 h in 4% paraformaldehyde and then overnight in 30% (w/vol) sucrose in PBS at 4°C. Tissues were mounted in OCT (Tissue-Tek), and 5 µm sections were cut and stored at –80°C.

### Immunofluorescent staining of kidney tissue

For immunostaining, sections were washed with Tris-buffered saline - 0.1% Tween 20 (TBSt). Blocking was performed with 10% (w/v) BSA diluted in TBSt for 30 min at room temperature and then incubated with primary and secondary antibodies diluted in TBSt with 10% (w/v) BSA overnight at 4°C and 1 h at room temperature, respectively. After washing with TBSt, tissue sections were mounted using VECTASHIELD Vibrance mounting medium (Vector Laboratories) to preserve fluorescence. Fluorescent images were acquired using a confocal microscope (Leica DMi8) or an Echo Revolve R4 microscope.

### Western blotting of kidney protein samples

Kidneys were homogenized with lysis buffer containing 250 mM sucrose, 10 mM triethanolamine, 50 mM sodium fluoride, 5 mM sodium pyrophosphate, 1 mM sodium orthovanadate, 10 mM 1,10-phenanthroline, and Complete protease inhibitor cocktail (Roche). Protein concentration was quantified by the bicinchoninic acid (BCA) protein assay (Pierce). Proteins were prepared in 1x Laemmli buffer and loaded into SDS-polyacrylamide gels, transferred to PVDF membranes, and blocked for 1 h with 10% (w/v) non-fat milk in TBST. Antibodies were diluted in 5% (w/v) non-fat milk in TBST. Incubation with primary and horseradish peroxidase-coupled secondary antibodies was performed overnight at 4°C and 1 h at room temperature, respectively. Enhanced chemiluminescence reagent was used for signal detection.

### Cell experiments and immunoblots

HEK293 (ATCC® CRL-1573) cells were transiently transfected with expression plasmids (see Table S1). Cells were grown at 37°C, with 5% CO2, in DMEM (Gibco) with 10% FBS, to a 70-80 % confluence and then transfected with Lipofectamine 2000 (Life Technologies). Forty eight hours after transfection, cells were lysed with a lysis buffer containing 50 mM Tris·HCl (pH 7.5), 1 mM EGTA, 1 mM EDTA, 50 mM sodium fluoride, 5 mM sodium pyrophosphate, 1 mM sodium orthovanadate, 1% (w/v) Nonidet P-40, 270 mM sucrose, and protease inhibitors (Complete tablets; Roche Applied Science and 10 mM 1,10-phenanthroline). Protein concentration was quantified using BCA (Biorad) assay and Western blots were performed as described above. For live microscopy studies, cells were seeded in glass bottom culture dishes (Nest, 801002) and transfection was performed as explained above. Constructs with fused fluorescent tags were used. Pictures of cells were taken 24-48 h post transfection.

For some experiments, simultaneous observation of multiple transfected proteins was achieved through observation of fluorescent label-tagged proteins and immunostaining of other proteins. For immunostaining in cells, twenty-four hours after transient transfection, cells were treated with PBS++ (containing 0.9 mM CaCl2 and 0.5 mM MgCl2) and subsequently fixed with 2% paraformaldehyde (PFA) in PBS++ for 15 minutes. Following fixation, cells were washed twice with PBS++. Next, they were incubated with a blocking solution (PBS++ containing 10% w/v BSA and 0.3% Triton X-100) for 1 hour at room temperature. The primary antibody was diluted in the blocking solution and incubated with the cells at 4°C overnight. After three washes with PBS++ containing 0.2% Tween, cells were incubated in the dark with mouse anti-rabbit Dylight 405 nm antibody (diluted 1:150 in blocking solution) for 1 hour at room temperature. Following three additional washes with PBS++ containing 0.2% Tween, mounting media (Vectashield Vibrance in PBS++, 1:20) was added to the cells.

### Immunoprecipitation

Immunoprecipitation of recombinant FLAG-tagged proteins was performed with FLAG M2 Magnetic Beads (Sigma) following the manufacturer’s instructions. Briefly, 1 mg of protein extract was prepared and diluted with immunoprecipitation buffer (1x TBS, 1% Triton-X-100, 1 mM EDTA (pH 7.5), 1 mM sodium orthovanadate, 10 mM sodium pyrophosphate, and 50 mM Sodium fluoride). 30 μl of Magnetic FLAG beads were added and incubated at 4°C for 2 h, after which they were washed three times with 500 μl of 1x TBS buffer with 0.05 % Tween. Bound proteins were then eluted with a denaturing buffer containing SDS, glycerol, bromophenol blue, and β-mercaptoethanol (2x Laemmli).

Immunoprecipitation of recombinant HA-tagged proteins was performed with HA Magnetic Beads (Pierce) following the manufacturer’s instructions. Briefly, 15 ul of anti-HA magnetic beads were washed three times with wash buffer (TBS + 0.1% tween) then 1 mg of lysate was added. The suspension was incubated at room temperature for 20 min. After incubation the beads were washed three times then eluted in Laemmli buffer.

### Generation of SLC12 knockout HEK293 cells

NKCC1 and KCCs were sequentially targeted for inactivation in HEK293 (ATCC, CRL-1573) cells. In a first step, guide CCGCTTCCGCGTGAACTTCG targeting *SLC12A2* at exon 1 was inserted into vector pX458 at the *Bbs*I site. HEK293 cells were then transfected with this targeting vector. Two days post transfection, cells were FACS sorted using EGFP fluorescence as a marker of vector expression, and single green cells were plated in 2-3 x 96-well plates. Clones were allowed to grow, transferred to 24-well plates, and tested for NKCC1 function. One clone lacking bumetanide-sensitive K^+^ influx (Fig. S7A) was selected and used for targeting K-Cl cotransport function. Note that under regular conditions, there is no KCC function detectable in HEK293 cells. The function can be elicited by exposing cells to the WNK inhibitor WNK463 (Fig. S7B, bars 1-3). Because transcript abundance of all four K-Cl cotransporters is similar in HEK293 cells (human protein Atlas, (*43*), we targeted the SLC12A4-7 genes at once. The following guides were selected: SLC12A4 (GGGCAGGTACACCCCCATGA), SLC12A5 (CGGCAGGTACACGCCCATGA), SLC12A6 (TGGGAGGTAGACACCC ATGA), SLC12A7 (CGGCAGGTAGACGCCGATGA). Cells were transfected with equal amounts of each pX458 vector, FACS sorted two days later, and clones were isolated and tested for function. Clone HEK293-ϕKCC-26 was selected and further characterized. As seen in Fig. S7B (bars 5 & 6), no activation of K^+^ influx is observed in these cells with WNK463, demonstrating absence of K-Cl cotransporter function.

### 83Rb uptake experiments

K^+^ influx measurements were performed using ^83^Rb as a tracer. Prior to the experiment, cells were plated in 24-well plates pre-coated with poly-L-lysine (0.1 mg/mL). They were then exposed to isosmotic or hyperosmotic (NKCC1 function) saline for 15 min preincubation in the presence or absence of 1 μM WNK463 (Sigma Aldrich). The isosmotic saline contained in mM: 140 NaCl, 5 KCl, 1 CaCl2, 0.8 MgSO4, 1 glucose, 10 HEPES, pH 7.4. The hyperosmotic saline was of identical composition but contained 80 mM sucrose. After aspiration, the preincubation solution was replaced with identical saline containing 100 μM ouabain, 0.25 μCi/mL ^83^Rb with or without 20 μM bumetanide (NKCC1 function), 1 μM WNK463 (to activate KCC function), 10 μM ML077 (VU0240551, to inhibit KCC function). Before the flux experiment started, two 5 μL aliquots of radioactive saline were sampled, added to vials containing 5 mL scintillation fluid, and used as standards. Each flux condition was measured in triplicate. The uptake was terminated after 15 min by aspiration of the radioactive solution and 3 rapid washes with ice cold saline. Cells were then lyzed with 500 μL NaOH 0.2N for 1 hour, then neutralized with 250 μL acetic acid glacial. Lysates (150 μL for scintillation counting and 20-30 for protein assays) were sampled. K^+^ influx was calculated and expressed in nanomole K^+^ per mg protein per min.

### Generation of a Long-WNK1 (L-WNK1) knockout HEK293 cell line

The PX458 polycistronic plasmid was used that allows for simultaneous SpCas9, GFP, and gRNA expression as described in the protocol by Ran et al. (*44*). Briefly, a sgRNA targeting exon 1 of WNK1 (sequence TCCAGCGAACCGACCATGTC), selected with the Chopchop web tool for gRNA specificity and efficiency maximization (https://academic.oup.com/nar/article/47/W1/W171/5491735), was cloned into the PX458 plasmid. This was confirmed by Sanger sequencing. HEK293 cells were transfected with this plasmid as described in a previous section, detached 48 hours later, and subjected to FACS based on GFP fluorescence for single cell isolation into a 96-well plate. After clonal expansion, cells were subjected to Western Blot to assess the levels of WNK1 abundance and kinase activity by the measurement of pSPAK-S371 phosphorylation (the S motif site, previously identified as S373; see Q9UEW8 in Uniprot). The C3 clone was selected as the knockout cell line, as undetectable levels of WNK1 were found, as well as clearly reduced levels of SPAK phosphorylation (Fig. S10).

### Plasmids

All modified constructs (e.g. WNK4 mutants, FKBP12 and FRB fusion proteins, tagged proteins, etc.) were generated by Fast Cloning (*45*). Fast cloning was performed using the Phusion-Plus high-fidelity DNA polymerase and confirmed by Sanger sequencing or whole plasmid sequencing (Plasmidsaurus, USA). For plasmid information see Table S1. pSpCas9(BB)-2A-GFP (PX458) was a gift from Feng Zhang (Addgene plasmid # 48138 ; http://n2t.net/addgene:48138 ; RRID:Addgene_48138)

### Antibodies

Antibody sources, dilutions, and validation data and references are provided in the Table S2 and Fig. S9.

### Statistical analysis

All statistical analyses were performed using [R Version 2024.04.2+764]. For comparisons between two independent groups, Student’s t-test was applied to evaluate statistical significance. Data are presented as mean ± standard deviation, and statistical significance was defined as p < 0.05 for all tests. For comparisons among multiple groups, one-way analysis of variance (ANOVA) tests were conducted followed by Tukey’s post hoc tests.

## Acknowledgements

We thank David Adams (Sanger Institute) for gifting us the NRBP1-Flag construct, David Pearce (UCSF) for providing a TSC22D3 construct, and Alejandro Rodríguez Gama (Northwestern Univ.) for providing the TSC22D-BFP construct. We thank Mariela Guadalupe Contreras Escamilla, Berenice Díaz Ramos, Marysol González Yáñez, Tania Perez Benhumea, and Anahi Leticia Aguilar Lopez, from the animal facility, for the help with the mouse colonies. We thank the Instituto Nacional de Cancerología - Advanced Microscopy Applications Unit (ADMiRA), RAI, UNAM (RRID:SCR_022788) and the Pathology department of the Instituto Nacional de Ciencias Médicas y Nutrición Salvador Zubirán for providing access to their microscopy equipment.

## Author contributions

Conceptualization: GMA, HCC, GG, MCB

Methodology: GMA, ED, MCB

Investigation: GMA, HCC, RA, TD, MTS, KGA, NV, ERO, ED, MCB

Formal analysis: GMA

Resources: BMC, ALS, NV, ED, DA, GG

Supervision: DHE, DRA, GG, MCB

Writing—original draft: GMA, MCB

Writing—review & editing: GMA, HCC, RA, ED, DE, DA, GG, MCB

Funding acquisition: ED, DE, DA, GG, MCB

## Data and materials availability

All data are available in the main text or the supplementary materials.

