## Supplementary material for "NRBP1 and TSC22D proteins impact distal convoluted tubule physiology through modulation of the WNK pathway"

Germán Magaña-Ávila *et al.*

**This PDF file includes:**

**Fig. S1.** TSC22D2-positive condensates are also observed in sporadic NCC-negative cells.
**Fig. S2.** NRBP1 acts in concert with long TSC22D isoforms to promote WNK-mediated SPAK phosphorylation.
**Fig. S3.** NRBP2 also promotes WNK-SPAK pathway activation.
**Fig. S4.** Comparison of the effect on SPAK phosphorylation of increased abundance of NRBP1 vs. co-expression of NRBP1 and TSC22D2.
**Fig. S5.** NRBP1 positive tubules in DCT-specific knockout mice.
**Fig. S6.** WNK4-CCTL1,2 mutant does not co-localize with TSC22D2 in cytoplasmic condensates in the absence of SPAK.
**Fig. S7.** Validation of SLC12 knockout cells.
**Fig. S8.** NRBP1 binds SALL3 in vitro.
**Fig. S9.** Validation of the TSC22D2 and NRBP1 antibodies used.
**Fig S10.** Generation and validation of an L-WNK1-knockout HEK293 cell line.
**Table S1.** Plasmids
**Table S2.** Antibodies

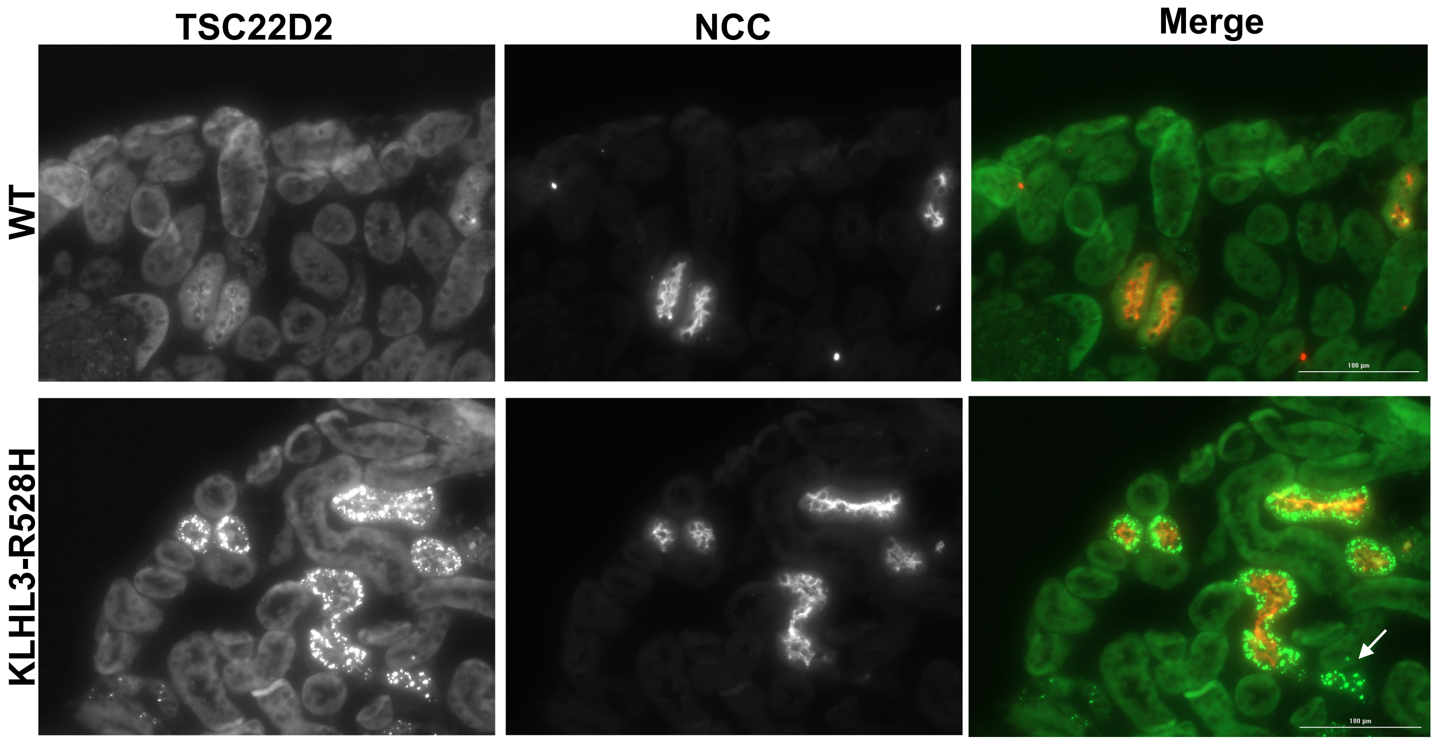


**Fig. S1. TSC22D2-positive condensates are also observed in sporadic NCC-negative cells.** Kidney tissue slices from wild type and KLHL3-R585H homozygous knockin mice were co-stained with antibodies against NCC and TSC22D2. With the TSC22D2 antibody, condensates were mainly observed in the cytoplasm of DCT cells from KLHL3-R528H knockin mice, but also in a few NCC-negative tubules. The white arrow indicates an example of such a tubule.

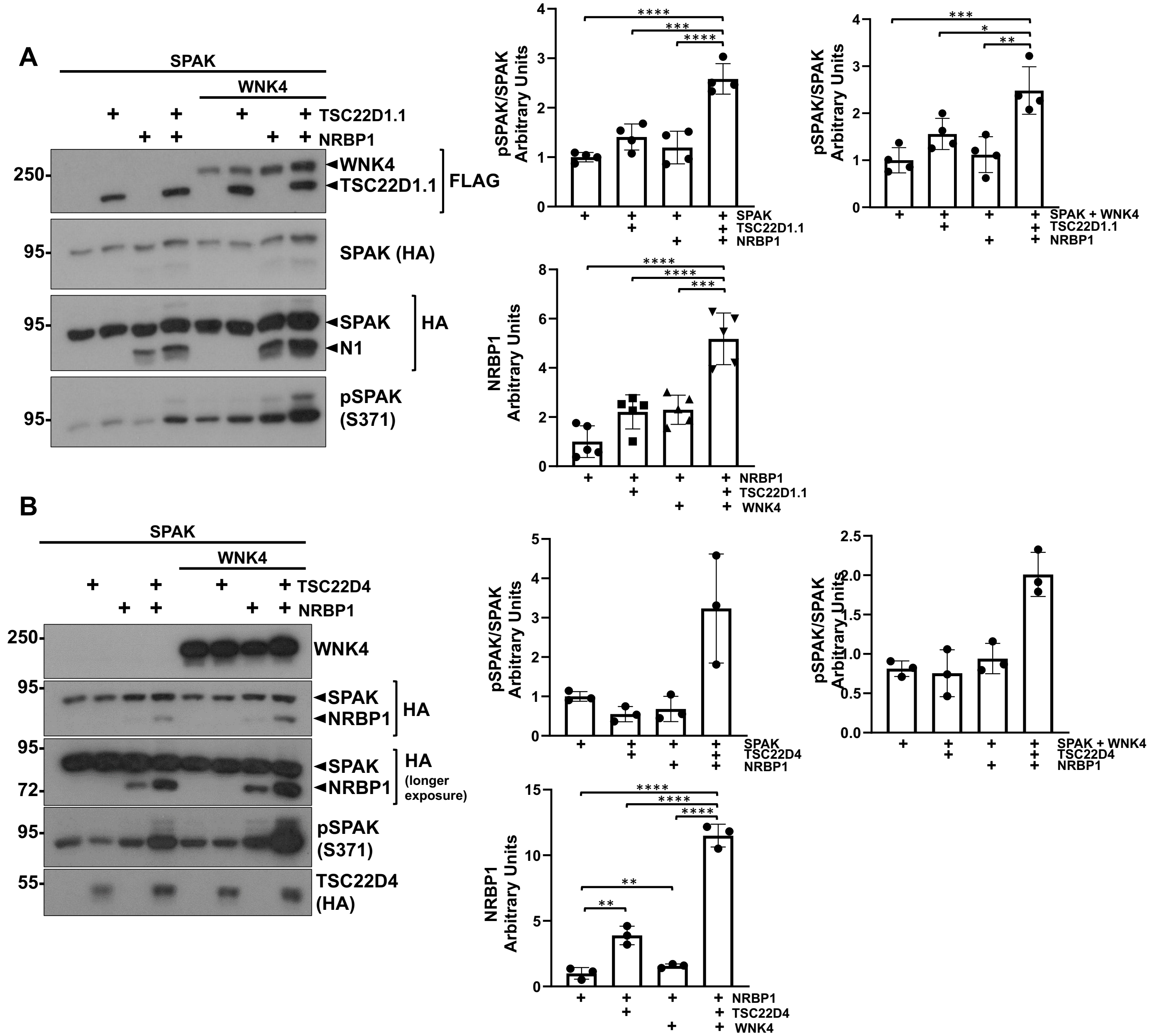


**Fig. S2. NRBP1 acts in concert with long TSC22D isoforms to promote WNK-mediated SPAK phosphorylation**. The effect of co-expression of NRBP1 with TSC22D1.1 **(A)** or TSC22D4 **(B)** on endogenous WNK's and WNK4´s ability to phosphorylate SPAK was assessed in HEK293 cells. Cells were transiently transfected with SPAK, WNK4, NRBP1 and the different TSC22D long isoforms as indicated. 48 hours post transfection immunoblots were performed to confirm expression of transfected proteins and to analyze phosphorylation levels of SPAK. An increase in SPAK phosphorylation levels was observed upon co-expression of NRBP1 and TSC22D1.1 or TSC22D4 in the absence of overexpressed WNK, perhaps due to an effect on the endogenous WNK. In the presence of WNK4, SPAK phosphorylation increased upon co-expression of NRBP1 with both long TSC22D proteins. Results of quantitation are shown in the graphs to the right. ANOVA followed by Tukey post hoc tests were performed to identify statistically significant differences. *$p<0.05$, **$p<0.01$, ***$p<0.001$, ****$p<0.0001$. At least three independent experiments were performed with each TSC22D isoform.

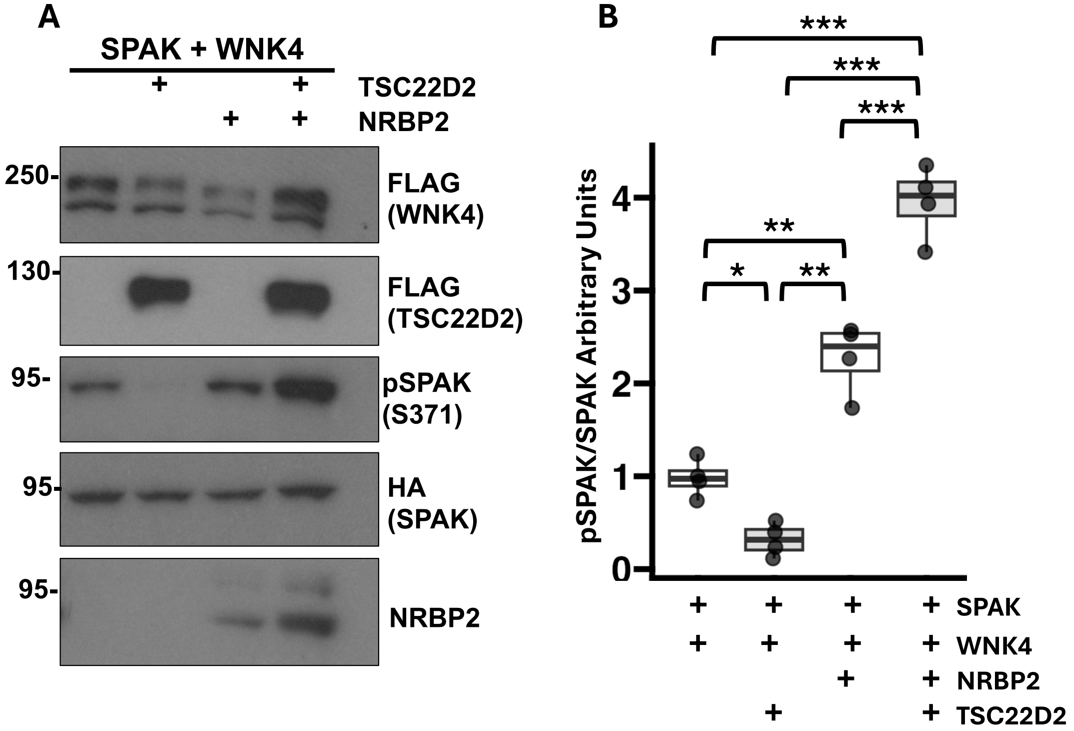


**Fig. S3. NRBP2 also promotes WNK-SPAK pathway activation.** (A) The effect of NRBP2 on WNK4-mediated SPAK phosphorylation was assessed in the absence and presence of TSC22D2. Cells were transiently transfected with SPAK, WNK4, NRBP2 and TSC22D2, as indicated. Co-expression of NRBP2 with TSC22D2 promoted an increase in the levels of pSPAK. Results of quantitation are shown in the graphs to the right. ANOVA followed by Tukey post hoc tests were performed to identify statistically significant differences. *$p<0.05$, **$p<0.01$, ***$p<0.001$. At least three independent experiments were performed.

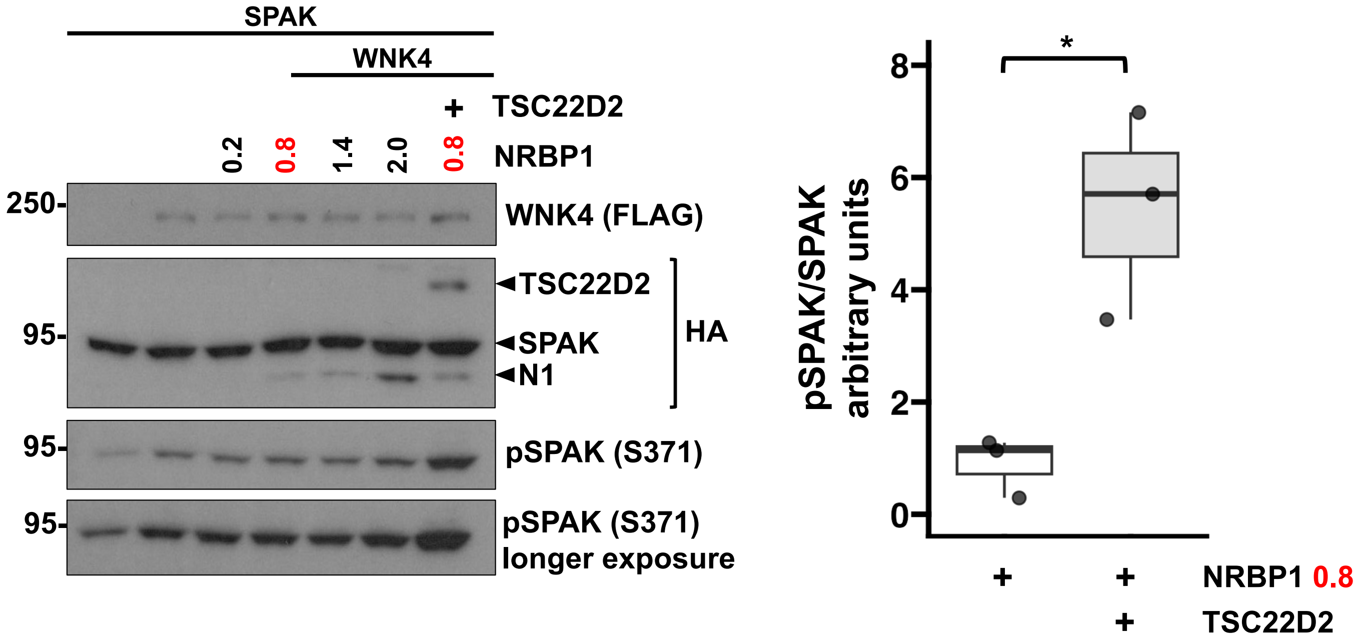


**Fig. S4. Comparison of the effect on SPAK phosphorylation of increased abundance of NRBP1 vs. co-expression of NRBP1 and TSC22D2.** HEK293 cells were transfected with SPAK, WNK4, and increasing amounts of NRBP1 as indicated. In addition, one group of cells was transfected with WNK4, SPAK, 0.8 mg of NRBP1 DNA, and TSC22D2. Despite observing a higher abundance of NRBP1 in the group transfected with the highest amount of NRBP1 alone than in the group with NRBP1 + TSC22D2, a clearly greater pSPAK signal was observed in the latter group. This suggest that the additive effect on SPAK phosphorylation of NRBP1 and TSC22D2 co-expression was not due to the effect of TSC22D2 on NRBP1 abundance. At least three independent experiments were performed with similar results. Student t test was performed to compare groups transfected with 0.8 μg of NRBP1 expression plasmid; *$p<0.05$.

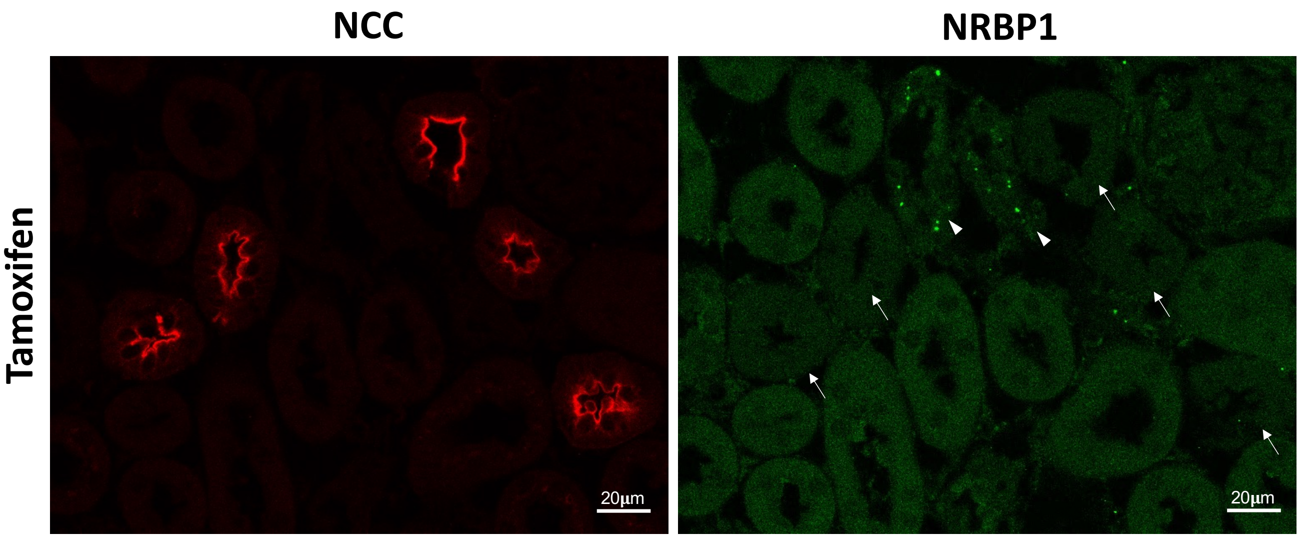


**Fig. S5. NRBP1 positive tubules in DCT-specific knockout mice.** Immunofluorescent staining of kidney sections from DCT-specific NRBP1 knockout mice. NRBP1-positive condensates were observed in sporadic cells that were NCC-negative (arrowheads), supporting the specificity of the cell type-specific targeting strategy and confirming that NRBP1-positive condensates are present in certain non-DCT cells. Also noticeable in this image is that NCC positive tubules (arrows) have no NRBP1-positive condensates and have a less intense diffuse cytoplasmic staining than surrounding tubules.

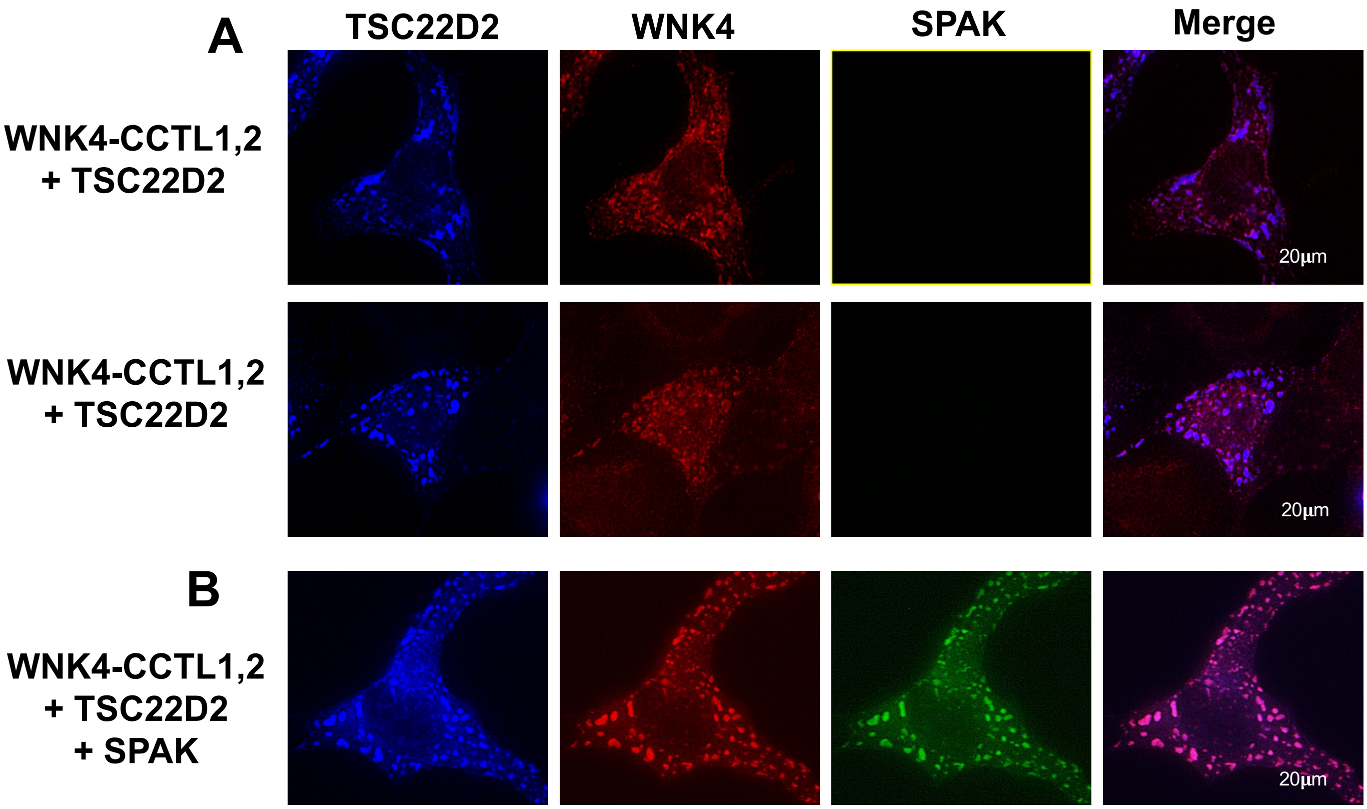


**Fig. S6. WNK4-CCTL1,2 mutant does not co-localize with TSC22D2 in cytoplasmic condensates in the absence of SPAK.** COS7 cells were transiently transfected with the WNK4-CCTL1,2 mutant. TSC22D2-BFP was co-transfected in the absence **(A)** or presence **(B)** of SPAK-GFP. Panel A shows TSC22D2- condensates surrounded by WNK4 condensates, with no signal overlapping. Panel B shows that in the presence of SPAK, TSC22D2 and the WNK4-CCTL1,2 mutant colocalize in condensates.

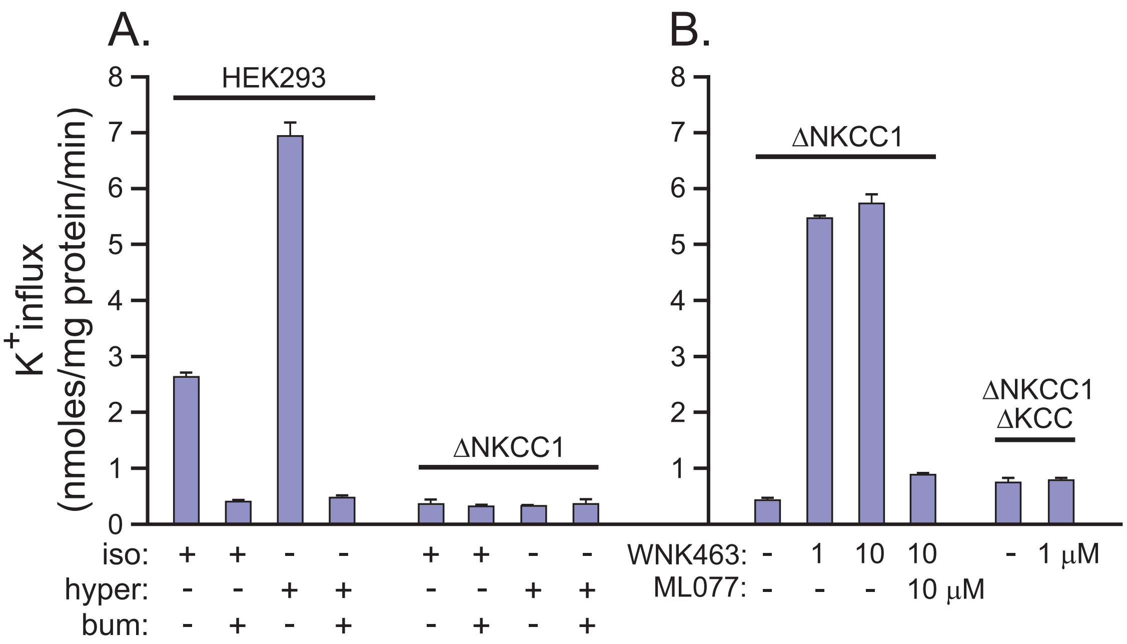


**Fig. S7. Validation of *SLC12* knockout cells. (A)** $^{83}$Rb uptake experiments were performed in wild type HEK293 cells and cells in which the *SLC12A2* gene was targeted by CRISPR-CAS9 (ΔNKCC1 cells). Experiments were performed under isosmotic and hyperosmotic conditions and in the absence or presence of the NKCC inhibitor bumetanide. $K^+$ influx was calculated and expressed in nanomole K+ per mg protein per min. **(B)** $^{83}$Rb uptake experiments were performed in ΔNKCC1 cells previously validated in (A) and in ΔNKCC DKCC cells that were generated by simultaneously targeting all KCC-encoding genes in the ΔNKCC1 cells. KCC activity was stimulated by preincubation with the WNK inhibitor WNK463. Uptake experiments were performed in the absence or presence of the KCC inhibitor ML077.

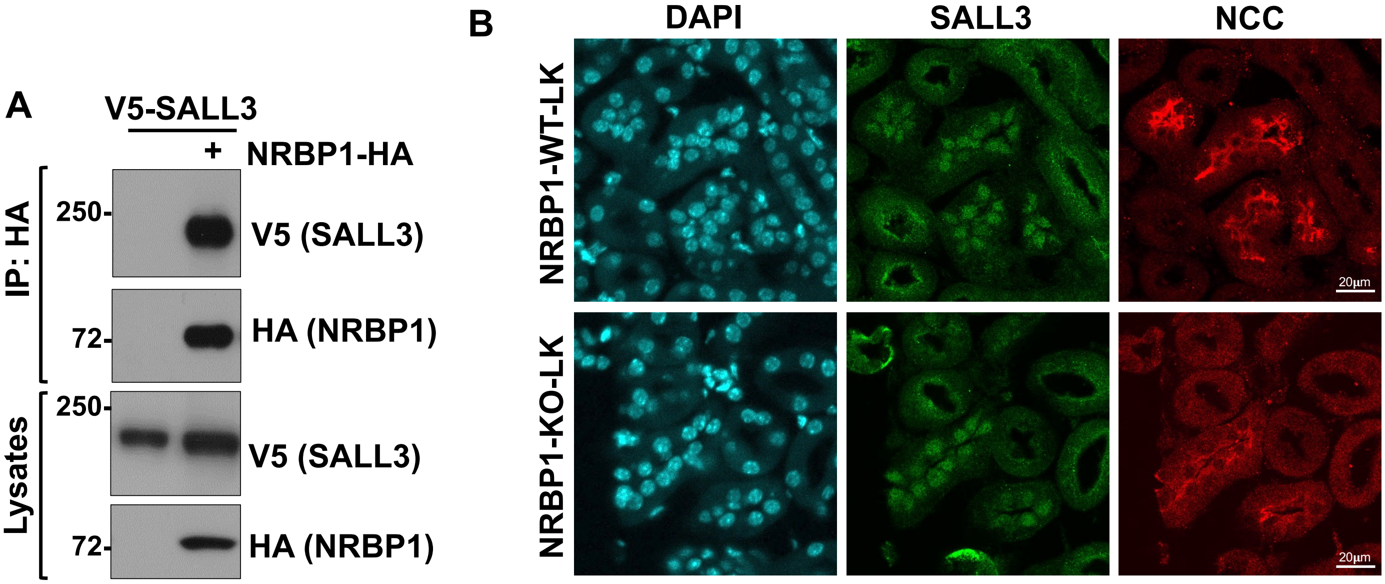


**Figure S8. NRBP1 binds SALL3 in vitro. (A)** HEK293 cells were transfected with V5-tagged SALL3 and HA-tagged NRBP1. HA-IP was performed to assess SALL3 co-immunoprecipitation. At least three independent experiments were performed with similar results. **(B)** In kidney sections stained with florescent probe-tagged antibodies a nuclear signal for SALL3 was only observed in DCT cells as previously reported (1). SALL3 nuclear localization in NRBP1 knockout mice was similar to that observed in control mice.

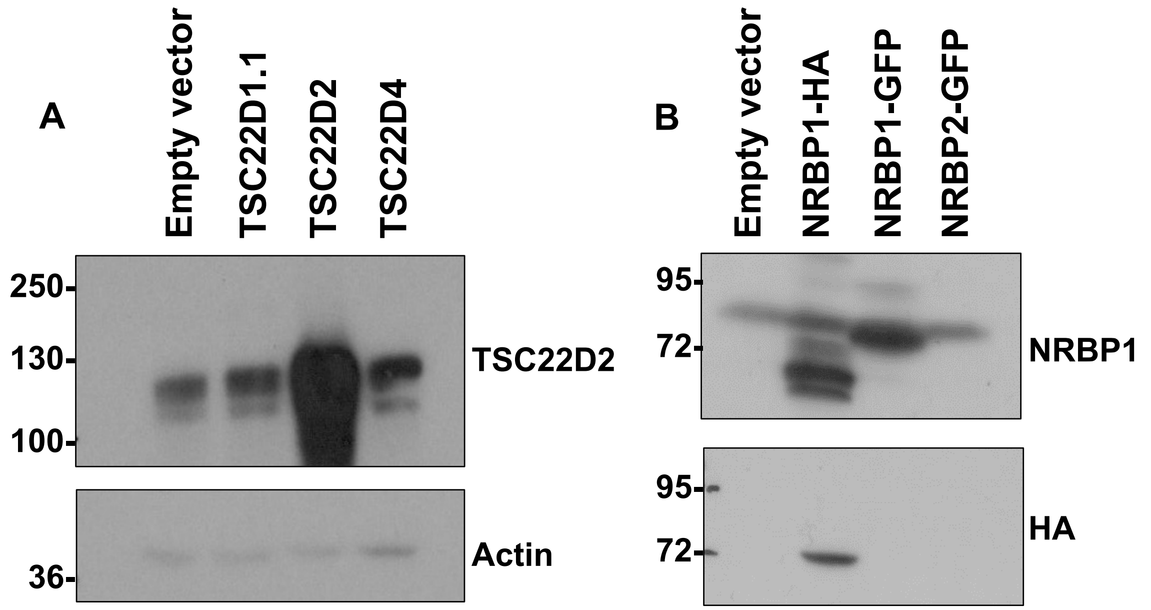


**Figure S9. Validation of the TSC22D2 and NRBP1 antibodies.** HEK293 cells were transfected with the indicated constructs and protein extracts were used to perform immunoblots with the TSC22D2 antibody **(A)** or the NRBP1 antibody **(B)**. Results show that the TSC22D2 antibody does not recognize other expressed TSC22D proteins and that the NRBP1 antibody does not recognize NRBP2. Immunofluorescence data presented in Fig. 5 also supports the specificity of this latter antibody.

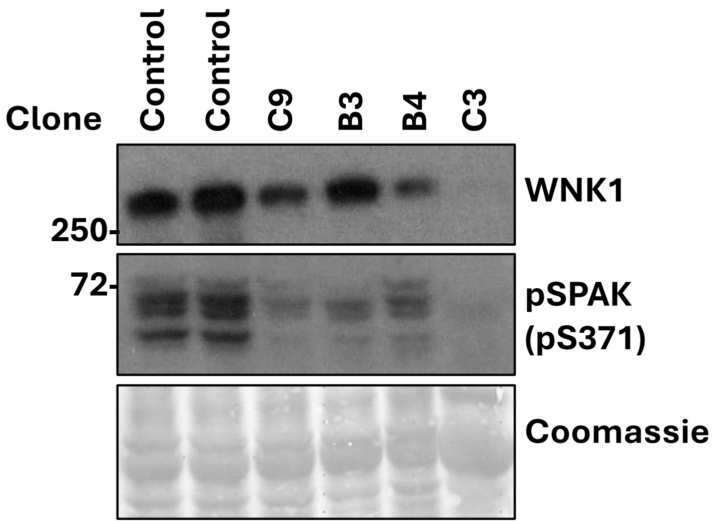


**Figure S10. Generation and validation of an L-WNK1-knockout HEK293 cell line.** An L-WNK1-knockout HEK293 cell line was generated with CRISPR-Cas9. Control lanes represent parental HEK293 cells. Individual clones, named after the well where they were originally grown, were assessed by Western blot. The C3 clone was identified as the L-WNK1 knockout, displaying undetectable WNK1 protein levels and significantly reduced pSPAK levels.

**Table S1. Plasmid constructs used**

| Plasmids | Name | Accession | Mutations | Tag | Species | Origin, ID | Ref. |
|---|---|---|---|---|---|---|---|
| **pcDNA5D FRT/TO HA NRBP1** | HA-NRBP1 | NP_001308286 | | HA | Human | MRC-PPU-Dundee, DU68362 | (*12*) |
| **pCMV5D HA TSC22D2** | HA TSC22D2 | NP_055594 | | HA | Human | MRC-PPU-Dundee, DU68295 | (*12*) |
| **pcDNA5D FRT/TO FLAG TSC22D2** | FLAG TSC22D2 | NP_055594 | | FLAG | Human | MRC-PPU-Dundee, DU80387 | (*12*) |
| **pCMV5D FLAG TSC22D1** | FLAG TSC22D1 | NP_904358 | | FLAG | Human | MRC-PPU-Dundee, DU68673 | (*12*) |
| **pcDNA5D FRT/TO FLAG TSC22D4** | FLAG TSC22D4 | NP_001289972 | | FLAG | Human | MRC-PPU-Dundee, DU77694 | (*12*) |
| **pCMV5D FLAG TSC22D3** | FLAG TSC22D3 | NP_001305397 | | FLAG | Human | MRC-PPU-Dundee, DU72786 | This study |
| **GFP-HA-SPAK-pEGFPC1** | GFP-HA-SPAK | AF099989.1 | | GFP HA | Human | MRC-PPU-Dundee, DU6188 | (*12*) |
| **pcDNA 3.1 (-) WNK4 FLAG** | WNK4 FLAG | NP_115763 | | FLAG | Human | MCB Lab | This study |
| **pcDNA 3.1 (-) WNK4 HA** | WNK4 HA | AAO21955 | | HA | Mouse | Lifton Lab | (*52*) |
| **pcDNA 3.1 (-) WNK4 CCTL1 FLAG** | WNK4 CCTL1 FLAG | | (F476A, F478A) | FLAG | Human | MCB Lab | This study |
| **pcDNA 3.1 (-) WNK4 CCTL2 FLAG** | WNK4 CCTL2 FLAG | | (V701A, V703A) | FLAG | Human | MCB Lab | This study |
| **pcDNA 3.1 (-) WNK4 CCTL1,2 FLAG** | WNK4 CCTL1 CCTL2 FLAG | | (F476A, F478A) , (V701A, V703A) | FLAG | Human | MCB Lab | This study |
| **pcDNA 3.1 (-) WNK4 RFAA FLAG** | WNK4 RFAA FLAG | | (R1016A, F1017A) | FLAG | Human | MCB Lab | This study |
| **pcDNA 3.1 (-) WNK4 CCTL1,2 RFAA (WNK4-TM) FLAG** | WNK4 CCTL1 CCTL2 RFAA FLAG | | (F476A, F478A) , (V701A, V703A), (R1016A, F1017A) | FLAG | Human | MCB Lab | This study |
| **mCherry-HA-SPAK** | mCherry-HA-SPAK | | | HA | | MCB Lab | This study |
| **pcDNA5D FRT/TO GFP TSC22D2** | EGFP TSC22D2 | NP_055594 | | | | MRC-PPU-Dundee, DU10054 | This study |
| **pcDNA5D FRT/TO BFP TSC22D2** | BFP TSC22D2 | NP_055594 | | | | MCB Lab | This study |
| **pCMV5D EGFP V5 TSC22D3** | EGFP V5 TSC22D3 | NP_001305397 | | GFP V5 | Human | MCB Lab | This study |
| **pcDNA 3.1 (-) WNK4-TM-mCherry-FKBP12 FLAG** | WNK4-TM-FKBP12 | | (F476A, F478A) , (V701A, V703A), (R1016A, F1017A) | FLAG | Human | MCB Lab | This study |
| **pcDNA5D FRT/TO GFP-FRB-TSC22D2** | TSC22D2-FRB | NP_055594 | | | Human | MCB Lab | This study |
| **pSpCas9(BB)-2A-GFP (PX458)** | Cas9 2A-EGFP | | | 3x FLAG | S. pyogenes | Addgene #48138 (F. Zhang) | (*44*) |

**Table S2. Antibodies used**

| Antibody | Dilution factor | Source | ID | Validation Ref. |
|---|---|---|---|---|
| **Rabbit Anti-TSC22D1.1** | 1:1000 (WB) 1:200 (IF) | Bethyl | A303-581A | This paper (Fig 3) |
| **Sheep Anti-TSC22D2** | 3µg/ml (WB) 1:200 (IF) | MRC-PPU-Dundee | DA056 | This paper (Fig 3, S10) and (*12*) |
| **Sheep Anti-TSC22D2** | 3µg/ml (WB) 1:200 (IF) | MRC-PPU-Dundee | S952B | This paper (Fig 3, S10) and (*12*) |
| **Sheep Anti-NRBP1** | 3µg/ml (WB) 1:200 (IF) | MRC-PPU-Dundee | DA204 | This paper (Fig 6, S10) and (*12*) |
| **Rabbit Anti-NRBP1** | 1:1000 (WB) 1:200 (IF) | Sigma | HPA029527 | This paper and (*12*) |
| **Sheep Anti-NRBP2** | 3µg/ml (WB) 1:200 (IF) | MRC-PPU-Dundee | DA240 | This paper (Fig S3) |
| **Rabbit Anti-WNK1** | 1:2000 (WB) 1:1000 (IF) | Bethyl | A301-515A | (*23*) |
| **Rabbit Anti-WNK4** | 1:5000 (WB & IF) | Ellison Lab | | (*46*) |
| **Rabbit Anti-NCC** | 1:5000 (WB & IF) | Ellison Lab | | (*53*) |
| **Sheep Anti-NCC 3P** | 3µg/ml | MRC-PPU-Dundee | S908B | (*54*) |
| **Sheep Anti-WNK1 pS382** | 3µg/ml | MRC-PPU-Dundee | S099B | (*29*) |
| **Sheep Anti-pSPAK S373/ pOSR1 S325** | 2 µg/ml (WB) 1:100 (IF) | MRC-PPU-Dundee | S670B | (*24*) |
| **FLAG** | 1:5000 | Sigma | A8592 | |
| **HA** | 1:2000 (WB) | Sigma | H6533 | |
